# Human iPSC-derived Microglial Cells Integrated into Mouse Retina and Recapitulated Features of Endogenous Microglia

**DOI:** 10.1101/2023.07.31.550858

**Authors:** Wenxin Ma, Lian Zhao, Biying Xu, Robert N. Fariss, T. Michael Redmond, Jizhong Zou, Wai T. Wong, Wei Li

## Abstract

Microglia exhibit both maladaptive and adaptive roles in the pathogenesis of neurodegenerative diseases and have emerged as a cellular target for central nervous system (CNS) disorders, including those affecting the retina. Replacing maladaptive microglia, such as those impacted by aging or over-activation, with exogenous microglia that can enable adaptive functions has been proposed as a potential therapeutic strategy for neurodegenerative diseases. To investigate microglia replacement as an approach for retinal diseases, we first employed a protocol to efficiently generate human-induced pluripotent stem cells (hiPSC)-derived microglia in quantities sufficient for in vivo transplantation. These cells demonstrated expression of microglia-enriched genes and showed typical microglial functions such as LPS-induced responses and phagocytosis. We then performed xenotransplantation of these hiPSC-derived microglia into the subretinal space of adult mice whose endogenous retinal microglia have been pharmacologically depleted. Long-term analysis post-transplantation demonstrated that transplanted hiPSC-derived microglia successfully integrated into the neuroretina as ramified cells, occupying positions previously filled by the endogenous microglia and expressed microglia homeostatic markers such as P2ry12 and Tmem119. Further, these cells were found juxtaposed alongside residual endogenous murine microglia for up to eight months in the retina, indicating their ability to establish a stable homeostatic state *in vivo*. Following retinal pigment epithelial (RPE) cell injury, transplanted microglia demonstrated responses typical of endogenous microglia, including migration, proliferation, and phagocytosis. Our findings indicate the feasibility of microglial transplantation and integration in the retina and suggest that modulating microglia through replacement may be a therapeutic strategy for treating neurodegenerative retinal diseases.

## Introduction

Microglia are the innate immune cells of the central nervous system (CNS), including the retina, and play pivotal roles in neuronal (Puñal et al., 2019; Anderson et al., 2019; Huang et al., 2012) and vascular development (Checchin et al., 2006; Ritter et al., 2006; Kubota et al., 2009), normal synapse formation (Stevens et al., 2007; Schafer et al., 2012; Schafer et al., 2015; Hong et al., 2016), maintaining local homeostasis in the neural environment (Li et al., 2018; Colonna et al., 2017; Sierra et al., 2013) and the regulation of immune activity (Okunuki et al., 2019). Conversely, they are also implicated in driving pathologic progression in various retinal diseases, including age-related macular degeneration (AMD) (Combadière et al., 2007; Ma et al., 2009, 2012; Karlstetter et al., 2015), glaucoma (Bosco et al., 2019; Ramírez et al., 2020), diabetic retinopathy (Altmann C et al., 2018; Xu et al., 2017), and uveitis (Broderick et al. 2002; Okunuki et al., 2019; Zhou et al., 2020).

Under homeostatic conditions in the adult retina, microglial cells are predominantly distributed in the IPL and OPL and vigilantly survey environmental changes through dynamic surveying behavior in their ramified processes (Lee et al., 2008). Their presence and homeostatic function are crucial for maintaining normal retinal functions, including the maintenance of synaptic function and integrity (Wang et al., 2016). Under normal conditions, microglial cells sustain equilibrium in their endogenous numbers via slow self-renewal (Réu et al., 2017). However, with the onset of pathology, this homeostasis can be disrupted following microglia activation, migration, and proliferation. Microglia repopulation in the retina following perturbation is achieved through both the proliferation of endogenous microglia and the infiltration of peripheral monocytes (Ma et al., 2017; Huang et al., 2018; Zhang et al., 2018).

Studies of microglia cell repopulation have indicated that retinal resident microglia can sustain their population with the local microglial cell dividing and migration if any perturbations do not exceed the threshold of the recovery speed by local neighbor microglia. However, in cases of more severe retinal injury or infection that cause significant redistribution of endogenous microglia, the reestablishment of retinal microglial homeostasis will, in addition, involve peripheral monocytes that infiltrate into the retina to take up residence as macrophages. The ability of the retina to incorporate exogenous monocytic cells suggests that microglia cell replacement employing exogenously introduced microglia may be feasible and can exert therapeutic effects post-injury. Inhibiting retinal microglia over-activation has shown efficacy in animal models of retinal injury (Zhao et al., 2011; Karlstetter et al., 2015; Au et al., 2022) and potential signal in early-phase clinical trials (Cukras et al., 2012). These observations suggest that the depletion of maladaptive microglia in pathological contexts and their replacement with microglia that have a more homeostatic phenotype may constitute a potential therapeutic strategy.

Studies of microglia have largely been performed in rodent-derived models, largely due to the accessibility of various transgenic disease models. However, several studies have indicated that genetic and functional differences exist between murine and human microglia (Dawson et al., 2018; Friedman et al., 2018; Ueda et al., 2016). For instance, microglia-expressed genes CD33 and CR1, which have been associated with Alzheimer’s disease (AD) risk in genome-wide association studies (GWAS), lack reliable orthologues in mice (Hasselmann et al., 2020). Additionally, over half of the AD risk genes that are enriched in microglia demonstrate <70% sequence homology between humans and mice (Hasselmann et al., 2020). There are also significant differences in the levels of protein expression of some microglia-expressed complement factors and inflammatory cytokines between humans and mice (Galatro et al., 2017; Gosselin et al., 2017; Smith & Dragunow, 2014). As a result, microglia from murine models may not accurately represent those found in human conditions (Burns et al., 2015), limiting their translational potential.

To study microglia of human origin, some investigators have attempted to isolate microglial cells from human tissue. However, owing to limitations in sample availability, and the rapid transcriptomic changes that ex vivo microglia undergo post-isolation, these studies have been technically constrained (Bohlen et al., 2017; Butovsky et al., 2014; Gosselin et al., 2017). As an alternative to primary microglia, human-induced pluripotent stem cells (iPSCs) offer a wealth of possibilities and have been increasingly used to generate human microglia via differentiation in vitro (Muffat et al., 2016; Pandya et al., 2017; Abud et al., 2017; Douvaras et al., 2017; Haenseler et al., 2017; Takata et al., 2017). This approach has enabled the generation of a large quantity of cells of a specified genetic background, enabling the creation of in vivo human microglia cell models through xenotransplantation of hiPSC-derived microglia into the murine CNS (Abud et al., 2017; Svoboda et al., 2019; Parajuli et al., 2021; Xu et al., 2020; Chadarevian et al., 2023).

In this study, we adopted a previously published protocol appropriate for culturing hiPSC-derived microglial cells (Muffat et al., 2016; Pandya et al., 2017; Abud et al., 2017; Douvaras et al., 2017; Haenseler et al., 2017; Takata et al., 2017). We characterized microglia differentiation by examining the RNA and protein expression levels of microglia-enriched genes. We also assessed the inflammatory responses and phagocytic functions of hiPSC-derived microglia *in vitro*. We then established a human iPSC-derived microglia cell model through the xenotransplantation of human iPSC-derived microglial cells into the retina of an adult mouse. We found that when transplanted by subretinal injection, human iPSC-derived microglial cells were able to migrate into the retina where native retinal microglia reside and acquire a morphology resembling endogenous mouse microglia, and express microglia signature markers. These grafted cells persisted in the mouse retina for at least eight months and responded to RPE cell injury in ways resembling endogenous mouse microglia. Xenografting of hiPSC-derived microglia into mouse retina has the promise of being used to create in vivo models of retinal disease and injury to evaluate the preclinical efficacy of potential therapeutic agents, as well as to evaluate microglia transplantation itself as a potential therapeutic intervention.

## Results

### Differentiation and characterization of human iPSC-derived microglia

We used five distinct human iPSC lines for microglia cell differentiation, including the first-available iPSC line, KYOUDXR0109B, from ATCC, and four other lines (NCRM6, MS19-ES-H, ND2-AAVS1-iCAG-tdTomato, and NCRM5-AAVS1-CAG-EGFP, all from the National Heart, Lung, and Blood Institute (NHLBI), NIH). Our approach to microglia differentiation was informed by our previous work with primary mouse retinal microglia cell culture (Ma et al., 2009) and a variety of established microglia cell differentiation protocols (Muffat et al., 2016; Pandya et al., 2017; Abud et al., 2017; Douvaras et al., 2017; Haenseler et al., 2017; Takata et al., 2017). We opted for the myeloid progenitor/microglia cell floating culture method (Van et al., 2013; Haenseler et al., 2017) for its simplicity, efficiency, and consistency, which enables the generation of a large and uniform population of microglial cells.

The differentiation process involved three key stages: embryoid body formation, myeloid progenitor cell generation, and microglia cell maturation (Fig.1 A-D). Following myeloid differentiation, floating myeloid progenitor cells were harvested and allowed to differentiate further for two weeks in 6-well plates under conditions promoting microglial differentiation. We modified this step to include additional differentiation factors in the differentiation medium, including IL34, CSF1, CX3CL1, TGFB1, and TGFB2. We found that these factors promoted microglial morphological ramification and process elongation. Immunohistochemical analysis of the resulting cells showed that among CD34(+) cells, 98.6% were immunopositive for IBA1 (Fig.1E, F), and 98.5% were immunopositive for P2RY12 (Fig.1E, G). Immunostaining with myeloid cell markers CX3CR1, CD68, and CD11b showed positivity in 88%, 99.7%, and 94.3% respectively, demonstrating the high efficiency of differentiation achieved by this procedure (Fig.1. Suppl). Most of the resulting hiPSC-derived microglia showed spindle-shaped morphologies, with some displaying short ramifications in their processes (Fig.1E), resembling those observed in primary mouse retinal microglia cultures (Ma et al., 2009). Floating myeloid progenitor cell harvest could be performed repeatedly over three months following culture establishment, providing a steady and consistent supply for further microglia differentiation and generation.

**Figure 1.**
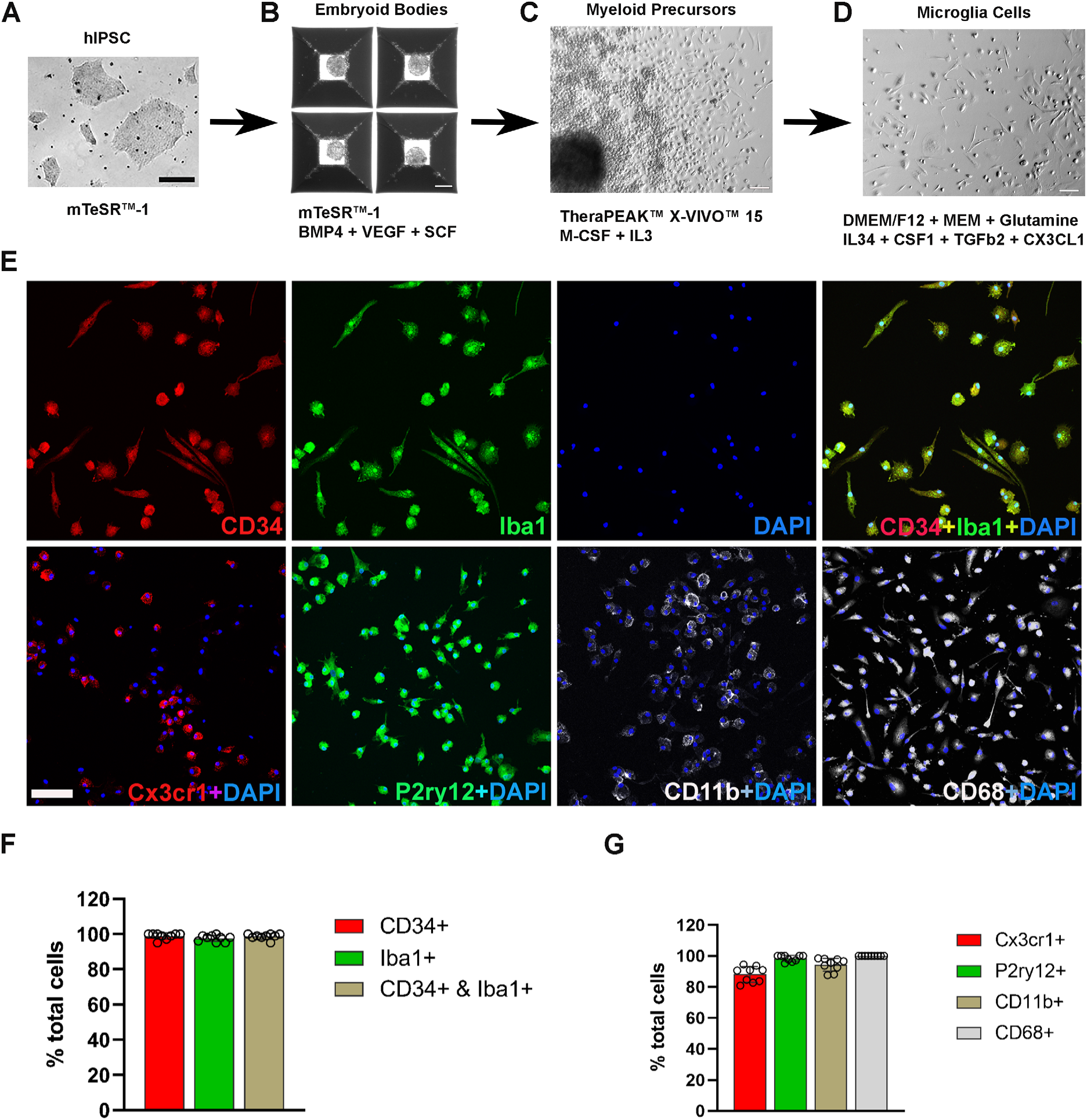
Differentiation and characterization of human iPSC-derived microglia. (A) Human iPSCs were cultured in a 6-well plate. Scale bar = 200µm. (B) Embryoid bodies formation was enabled in AggreWell™800 plate at day eight in culture medium mTeSR1 plus BMP4, VEGF, and SCF. Scale bar = 200µm. (C) Image of a myeloid precursor cluster following 1-month culture of embryoid bodies in TheraPEAK™ X-vivo™-15 Serum-free Hematopoietic Cell Medium with added M-CSF and IL3. Scale bar = 50µm. (D) Image of microglial cells in maturation culture for two weeks with DMEM/F12 plus nonessential amino acids, glutamine, IL34, CSF1, TGFb2, and CX3CL1. Scale bar = 50µm. (E) Immunohistochemical staining for Iba11 and human CD34, CX3CR1, P2RY12, CD11b, CD68. Scale bar = 100µm. (F) Cell counts and colocalization analysis of (F) CD34- and Iba11- positive cells and (G) positivity for myeloid cell markers CX3CR1, CD11b, activation marker CD68, and microglia marker P2ry12 in differentiated microglia.

A comparative RNAseq analysis between differentiated hiPSC-derived microglia collected at the end of the protocol vs. floating myeloid progenitor cells revealed a significant upregulation of microglia-enriched gene expression following microglial differentiation, including *CX3CR1, P2RY12, P2RY13, AIF1, TREM2, GPR34, CD53, CTSS, C3AR1*(Fig.2A-C). Moreover, hiPSC-derived microglia exhibited higher expression of genes associated with inflammation, apoptosis regulation, phagocytosis, lipid metabolism, and immune responses. The floating myeloid progenitor cells showed higher expression of hematopoietic/myeloid cell lineage genes (Fig. 2D). Ingenuity Pathway Analysis (IPA) of differentially expressed genes identified IL6, IL1β, and STAT3 as central signaling hubs critical to the regulation of inflammatory responses in microglia (Fig. 2E, Fig. 2 Suppl A, Suppl Table 3). We also compared the gene expression profiles between differentiated hiPSC-derived microglial cells vs. human microglia isolated ex vivo from the fetal and adult brain (Fig.2. Suppl. B, C and D; Suppl. table 1 and table 2)(Abud et al., 2017; Douvaras et al., 2017, Muffat et al., 2016, Böttcher et al., 2019, van der Poel et al., 2019, Eisenberg et al. 2013). The results of the correlation analysis indicated that hiPSC-derived microglial cells demonstrated an expression profile comparable to those in fetal and adult microglia in vivo but were more distinct from those in monocytes and inflammatory monocytes. Thus, our method of obtaining differentiated microglia can be employed to reliably and efficiently generate a large population of homogenous functional microglial cells of human origin.

**Figure 2.**
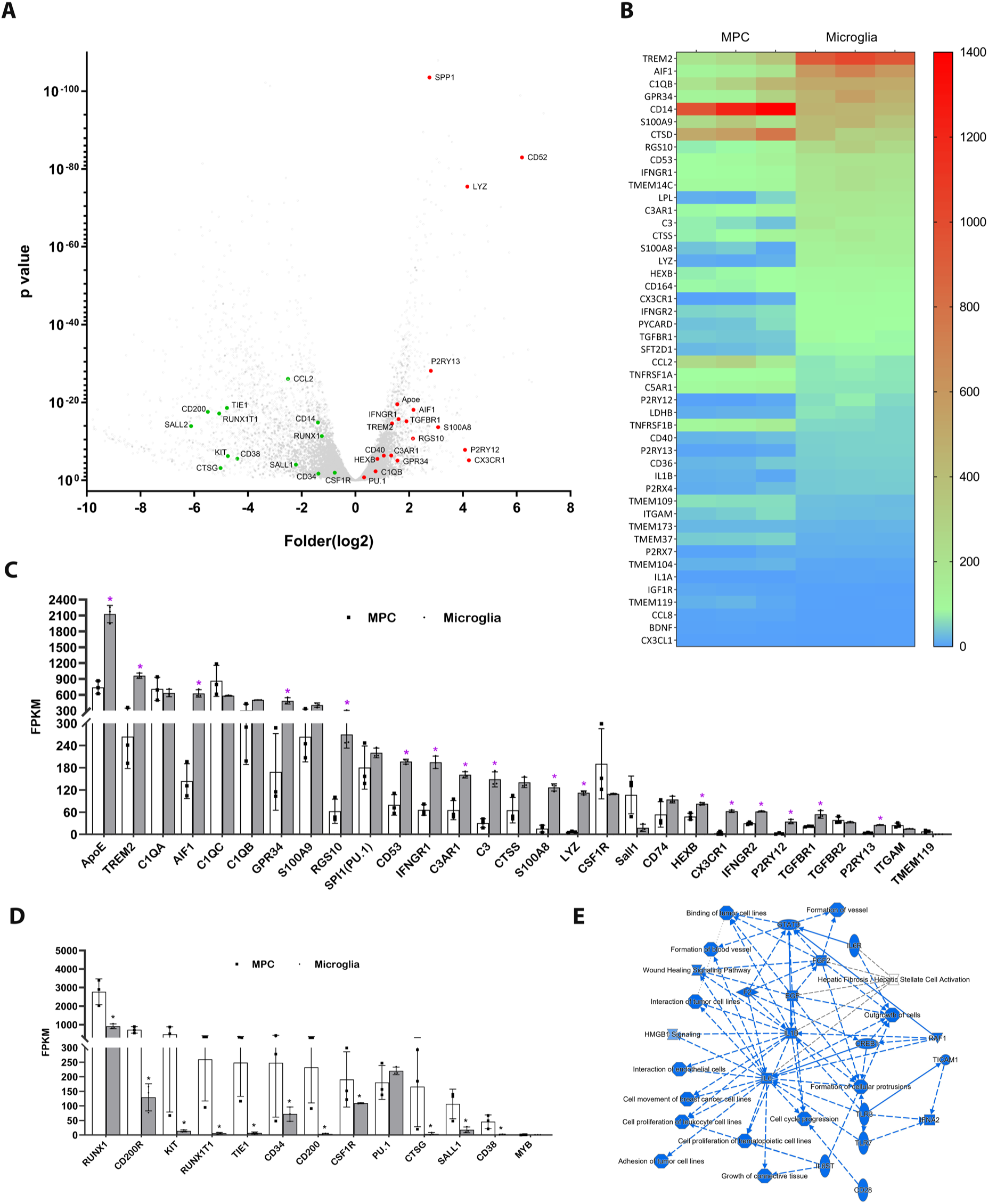
Profiling of genes differentially expression between differentiated microglial cells vs. myeloid progenitor cells (MPC) using bulk RNAseq analysis. (A) Volcano plot showing representative genes that were either upregulated (red) or downregulated (green) in differentiated microglia vs. MPCs. (B) Heat map showing increased expression of microglia-enriched genes in differentiated microglia vs. MPCs. (C) Histogram comparing the expression levels of microglia-enriched genes expression in terms of Fragments Per Kilobase of transcript per Million mapped reads (FPKM). * P<0.05. (D) Histogram comparing expression levels of myeloid cell lineage genes in human iPSC-derived MPC and microglia cells using FPKM. * P<0.05. (E) Graphic signaling pathway analysis with IPA analysis highlighting IL6 and IL1b as signaling hubs in differential gene expression patterns in differentiated microglia vs. MPCs.

## Human iPSC-derived microglia show inflammation responses and phagocytosis activity

Microglia play crucial roles in mediating inflammatory responses to stimuli and in phagocytosing pathogens. To investigate these functions further, we stimulated hiPSC-derived microglia with lipopolysaccharide (LPS) and analyzed their responses. Transcriptomic profiling using bulk RNAseq revealed that the primary responses to LPS stimulation involved changes in *IL6, IL1β, IL1α, TNFα,* and *IFNγ* signaling (Fig.3A, Suppl. Table 3), indicating the ability of hiPSC-derived microglia to demonstrate classical activation. This was confirmed through expression analysis using QRT-PCR and protein multiplex profiling, which showed a 50- to 800-fold increase in the expression of *IL6, IL1α, IL1β, TNFα, IL8, CXCL10,* and *CCL2* mRNA after 6 hours of LPS stimulation (Fig.3B). Similarly, we observed a significant increase in protein expression levels of these cytokines in cell lysate and culture medium (Fig.3C, D). We also treated hiPSC-derived microglia with IFNγ and a combination of IFNγ+LPS (Figure 3. Suppl.), and the results demonstrated that the combination of IFNγ+LPS promoted *IL1α, IL1β, TNFα, CCL8,* and *CXCL10* expression. These findings indicate that hiPSC-derived microglia, akin to microglia in vivo, exhibited strong responses to LPS and IFN*γ* stimulation.

**Figure 3.**
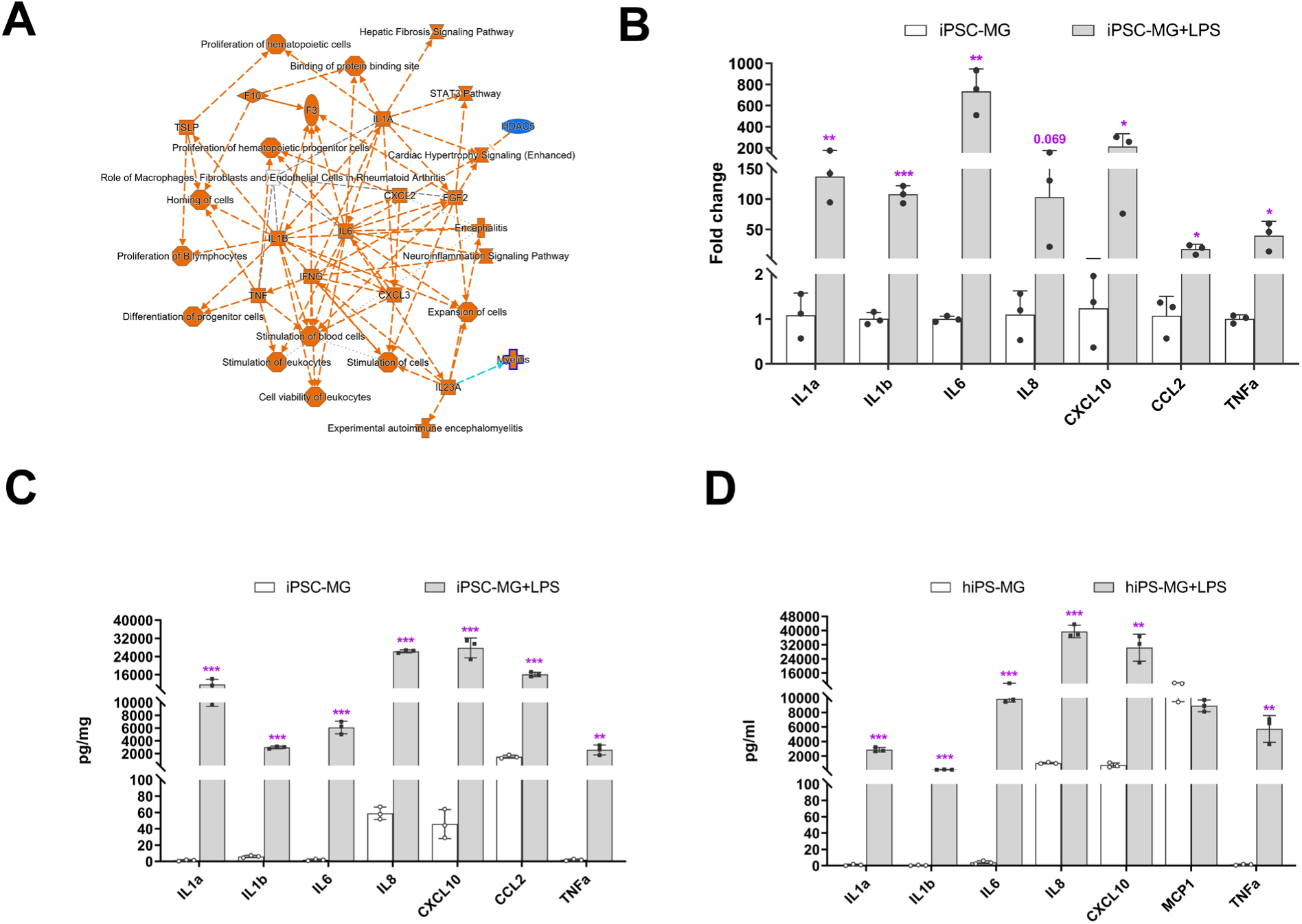
Inflammation responses of hiPSC-derived microglial cells following Lipopolysaccharide (LPS) stimulation. (A) Ingenuity Pathway Analysis (IPA) differentially-expressed genes (fold change >2-fold, p < 0.05) between LPS-treated and control hiPSC-derived microglial cells., demonstrating activation of core pathways involving IL6, IL1a, IL1b, and IFNG. (B) Assessment of mRNA expression of selected genes for inflammatory cytokines using QRT-PCR demonstrated increased expression following LPS (0.1µg/ml) stimulation for 6hrs. These changes corresponded to increases in the protein expression levels of inflammatory cytokines following 24 hours of LPS stimulation as measured with a Multiplex kit (Millipore) on celllysate (C) and in conditioned media (D). The data in (C) and (D) are presented as means ± SEM.* p<0.05, ** P<0.01, *** P<0.001, **** P<0.0001.

Microglia are local immune cells in the CNS, functioning as phagocytes involved in clearing apoptotic or necrotic cells, and cell debris (Green et al., 2016), remodeling neuronal connectivity by engulfing synapses, axonal and myelin debris (Paolicelli et al., 2011), and removing pathogens by direct phagocytosis (Nau et al., 2014). To assess the phagocytic capability of hiPSC-derived microglia, we exposed them to three different types of bioparticles: E. coli bacteria, zymosan, and bovine photoreceptor outer segments (POS) (Fig.4A-D). The cells altered their morphology in response and rapidly internalized the fluorescent-labeled particles (Fig. 4A-D, Fig.4. Suppl A and B). The engulfed bioparticles were condensed into perinuclear aggregates, likely within lysosomal bodies. They also demonstrated concurrent morphological changes into amoeboid-shaped cells (Fig.4D, E), resembling phenotypes demonstrated by retinal microglia phagocytosing photoreceptors in the context of photoreceptor degenerative pathologies in vivo (Zhao et al., 2015).

**Figure 4.**
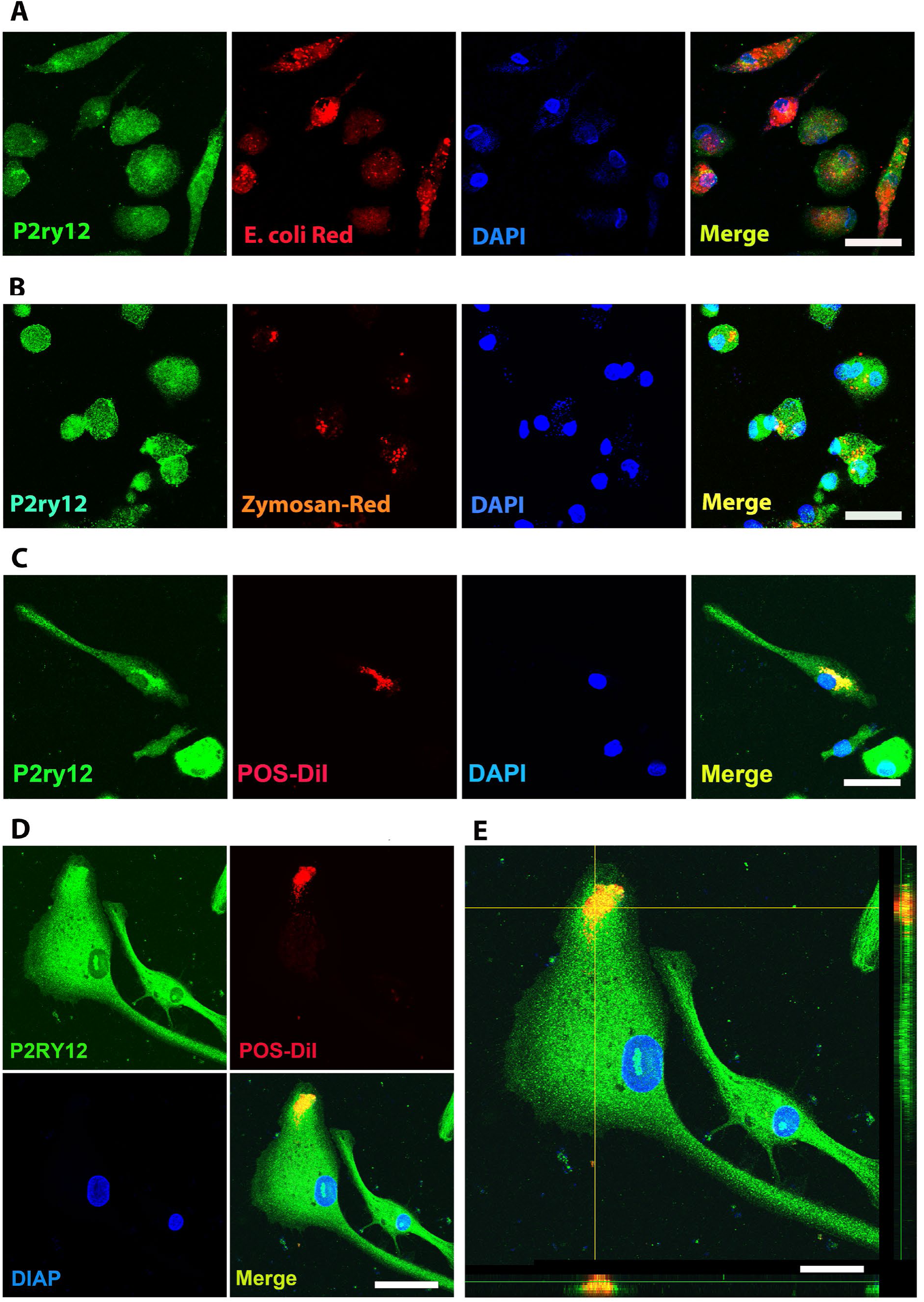
Human iPSC-derived microglia demonstrate robust phagocytosis. (A) pHrodo™ Red E. coli bioparticles were incubated in Human iPSC-derived microglia were incubated for 1 hour in either labelled pHrodo™ Red E. coli (A), pHrodo™ Red zymosan bioparticles (B), DiI-labeled bovine photoreceptor outer segments (POS) (C), and colabeled with anti-human P2RY12 antibody (green) and DAPI. Scale bar = 40µm. (D) A high-magnification view of a POS-containing intracellular vesicle within a labeled microglial cell is shown. Scale bar = 30µm.

### Transition of hiPSC-derived microglial cells to a homeostatic state within the mouse retina following xenotransplantation in vivo

To assess the ability of hiPSC-derived microglia to serve as microglia donor cell sources for transplantation, we conducted xenotransplantation experiments using humanized immunodeficient Rag2^-/-^;IL2rg^-/-^ ;hCSF1^+/+^ transgenic mice as recipients as previously employed (Svoboda et al., 2019; Xu et al., 2020; Chadarevian et al.,2023). Prior to xenotransplantation, recipient transgenic mice were pharmacologically depleted of endogenous retinal microglia by systemic administration of the CSF1R inhibitor PLX-5622 to create a depleted tissue niche for the integration of xenotransplanted microglia (Zhang et al., 2018). Two days following PLX-5622 treatment, adult transgenic mice were transplanted with 5000 hiPSC-derived microglia via injection into the subretinal space (Fig.5A). Tissue analysis at 4 and 8 months post-transplantation revealed that transplanted cells, which were marked by either tdTomato or EGFP expression, had migrated anteriorly from the subretinal space into the neural retina and were distributed across a wide retinal area within the inner and outer retinal layers including the ganglion cell layer (GCL), inner plexiform layer (IPL), and outer plexiform layer (OPL) (Fig.5B-D) in retinal loci typically occupied by endogenous microglia. Transplanted cells were immunopositive for IBA1, and human CD11b (hCD11b) (Fig.5H, I), as well as for microglia signature markers hP2RY12 and hTMEM119 (Fig.5D). Interestingly, the transplanted cells within the retina showed a ramified morphology typical of homeostatic microglia and demonstrated a regularly tiled “mosaic” distribution in their soma positions. These integrated cells were juxtaposed alongside residual endogenous murine microglia, which showed similar morphology and distribution as in endogenous conditions. This indicated that transplanted hiPSC-derived microglia responded to similar intraretinal cues and inter-microglia neighbor-neighbor interactions that guide the spatial and morphological organization of retinal microglia in vivo (Fig.5E-G). Similar observations were made for separate experiments involving the transplantation of tdtomato- and EGFP-expressing hiPSC-derived microglia (Fig.5B,C,H,I).

**Figure 5.**
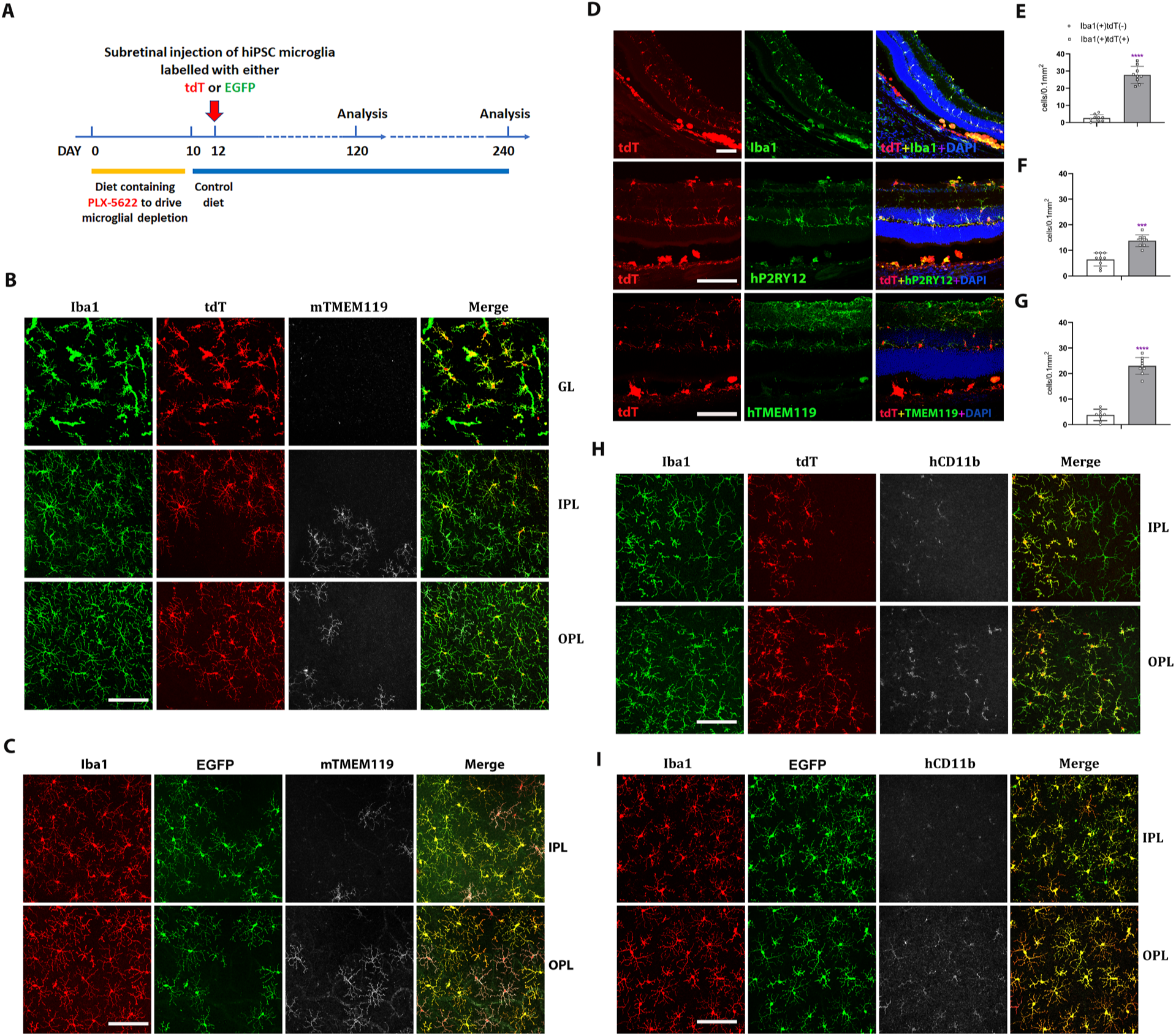
Human iPSC-derived microglial cells xenotransplanted into recipient mouse retina in vivo demonstrate recapitulation of endogenous distribution, cellular morphology, and stable integration for up to 4 months. (A) Schematic diagram shows the timeline for transplantation experiments. Two-month old adult transgenic Rag2^-/-^;IL2rg^-/-^;hCSF1^+/+^ mice were fed a PLX-5622-containing diet for 10 days before switching to standard chow. Two days following the resumption of standard chow, human iPSC-derived microglial cells expressing either tdTomato or EGFP were xenotransplanted into the subretinal space via subretinal injection (5000 cells in 1µl injection volume). Retinas were harvested for analysis 120 and 240 days following transplantation. (B) and (C) Retinas isolated post-transplantation were analyzed in flat-mounted tissue with confocal imaging. Transplanted hiPSC-derive microglia were visualized through their expression of tdtomato (TdT) (B) or EGFP (C), while endogenous mouse microglia were visualized using immunostaining for mouse Tmem119 (mTmem119). Imaging analysis was performed in separate layers of the retina including the ganglion cell layer (GL), inner plexiform layer (IPL), and outer plexiform layer (OPL). Scale bar = 100µm. (D) The retinal section showed human iPSC-derived microglial cells integrated into whole retinal layers (top panel) and positively stained with human P2ry12 and TMEM119 microglia signature markers. Scale bar = 100µm. The microglia cell number in GL, IPL, and OPL of host mouse retina were counted: mouse microglial cells (Iba11+, tdT-) and grafted human microglial cells (Iba11+,tdT+) were shown in (E), (F) and (G), respectively. *** P<0.001, **** P<0.0001. (H) and (I) showed tdT (H) or EGFP (I) labeled human iPSC-derived microglial cells in the IPL and OPL of the flat-mount retina with human CD11b staining. These results demonstrated that the infiltration of grafted hiPSC-derived microglial cells into the mouse retina is general in nature and not cell-line specific. Scale bar = 100µm.

We further evaluated the impact of human iPSC-derived microglia xenotransplantation on the host mouse retinal cells (Fig. 5 Suppl A-D). We examined the effect of microglia transplantation on endogenous Müller cell morphology and gliosis markers as retinal microglia have been described to interact with Müller glia to regulate overall retinal neuroinflammatory response (Wang et al., 2014). Immunostaining with glial fibrillary acidic protein (GFAP) and glutamine synthetase (GS) antibody (Fig. 5 Suppl A, B) indicated that the transplanted human iPSC-derived microglial cells did not trigger any upregulation of Müller cell gliosis markers or induced morphological changes or reactive proliferation for up to four months post-transplantation. Additionally, the laminar organization of the inner and outer retinal layers remained normal and similar to those in control retinas not subjected to transplantation (Fig. 5 Suppl. A-D), indicating that microglia transplantation had no adverse impact on the structural integrity of retina. Furthermore, immunostaining with the mouse CD11b antibody which marked the residual population of endogenous mouse microglia showed the spatial juxtaposition of these mouse microglia with transplanted human iPSC-derived microglia in a common mosaic of tiled cells (Fig. 5 Suppl. E, F), indicating the ability of transplanted hiPSC-derived microglia to integrate with and exchange signals with the residual endogenous microglial population, but not replaced.

### Transplanted hiPSC-derived microglia respond to induction of RPE cell injury with migration and proliferation

To monitor the longer-term consequences of microglia xenotransplantation, we extended analysis and examined the location and morphology of transplanted hiPSC-microglia up to 8 months post-transplantation. Analysis of retinal flat mounts confirmed that the tdTomato+ cells remained appropriately located in the GCL, IPL, and OPL, forming a mosaic-like distribution typical of endogenous microglia (Fig.6 Suppl A, B). These cells maintained expression of hP2RY12 and hTMEM119, markers of homeostatic retinal microglia, indicating their long-term integration within the recipient mouse retina (Fig.6 Suppl. A, B).

**Figure 6.**
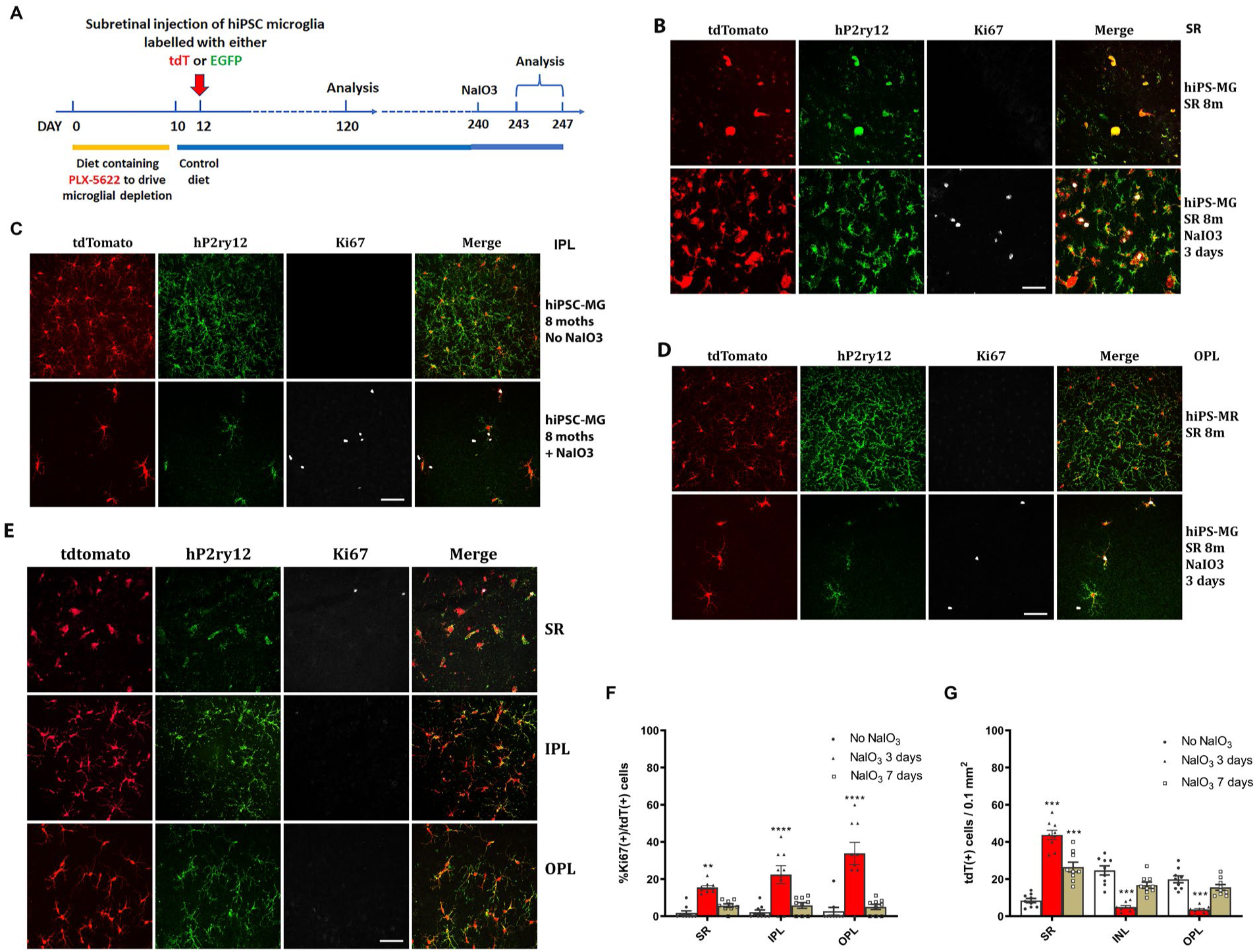
Migration and proliferation of hiPSC-derived microglia in the mouse retina after sodium iodate-induced RPE cell injury. (A) The schematic diagram shows the procedure of the experiment. After eight months post-transplantation of hiPSC-derived microglia, recipient animals were administered NaIO3 (30mg/kg body weight, intraperitoneal injection) to induce RPE injury. Retina were harvested at 3 and 7 days after NaIO3 administration and microglia numbers in the subretinal space monitored in retina and RPE-choroid flat mounts. (B) RPE-choroid flat-mounts demonstrate an increase of hiPSC-derived microglia (tdTomato+ and P2RY12+) in the subretinal space in response to RPE injury. A subset of subretinal microglia labelled for Ki67 indicating active proliferation. Scale bar = 60µm. (C) and (D) showed the number of P2ry12+&tdtomato+ human microglial cells in IPL (C) and OPL (D) decreased; some of them showed Ki67+ staining, Scale bar = 60µm. The cell count results showed in (F) and (G). (E) The retinal flat mount showed the number of P2ry12+&tdtomato+ human microglial cells in IPL and OPL that were repopulated, and the cells stopped dividing with lost the Ki67 staining at seven days after NaIO3 injection. The cell numbers were shown in (F) and (G). Scale bar = 60µm.

To evaluate the function of the transplanted hiPSC-derived microglia in the mouse retina and their ability to respond to injury in vivo, we subjected recipient mice 240 days post-transplantation to sodium iodate (NaIO3)-mediated RPE injury and analyzed retinal tissue 3 and 7 days post-injury (Fig 6A). In a previous study (Ma et al., 2015), we had characterized the responses of endogenous retinal microglia in this injury model; in the few days following injury, microglia within the neural retina migrated into the subretinal space, coming into close proximity to damaged RPE cells. This resulted in a transient decrease in microglia number in the IPL and OPL, which then recovered partially following the proliferation of the remaining microglia to replenish the depleted areas. We found that transplanted hiPSC-derived microglia demonstrated responses similar to endogenous microglia. Three days after NaIO3 injury, there was an increase in tdTomato+, hP2ry12+, hiPSC-derived microglia in the subretinal space, while their number decreased in the IPL and OPL (Fig.6B-D, G), indicating a migration of these cells from the neuroretina to the subretinal space. Some of the remaining tdTomato+ and P2ry12+ cells in the inner retina were positive for the cell-proliferation marker Ki67, indicating active cell division (Fig.6B-D, F). The numbers of Ki67+ tdTomato+ microglia peaked at three days post-injury and decreased thereafter (Fig.6B-F). Seven days post-injury, the numbers of tdTomato+ and P2ry12+ human iPSC-derived microglia increased in the IPL and OPL but decreased in the subretinal space (Fig.6E, G); Ki67+ tdTomato+ cell numbers also declined (Fig.6E, F). These findings suggest that once the hiPSC-derived microglia had replenished endogenous numbers in the inner retina, their division ceased. This response mirrors that of the endogenous mouse retinal microglia to NaIO3 injury (Ma et al., 2015).

Overall, these results demonstrate that the transplanted human iPSC-derived microglial cells retained a capacity for migration and proliferative responses to injury in a manner observed for endogenous microglia.

### hiPSC-derived microglial cells phagocytize debris or dead photoreceptor cells after NaIO3-induced RPE cell injury

Phagocytosis, a critical function of microglia, is essential both during development and in the resolution of pathological processes. Retinal microglia adaptively phagocytose and clear apoptotic photoreceptors in the rd10 mouse model of photoreceptor degeneration (Silverman et al., 2019). We found here that three days following NaIO_3_-induced RPE cell injury, tdTomato+ transplanted hiPSC-derived microglia migrated not only to the subretinal space but also into the photoreceptor layer (Fig.7A, C), coincident with the time of photoreceptor degeneration, when photoreceptor morphology becomes disrupted and photoreceptor density decreases (Fig.7A). Within the photoreceptor layer, hiPSC-derived microglia were observed to phagocytose photoreceptors as evidenced by the accumulation of intracellular autofluorescent material in their soma (Fig.7Ag, B, D), and their transformation into larger amoeboid cells (Fig. 7B, yellow triangle) containing arrestin-immunopositive photoreceptor-derived debris. mRNA analysis for human-specific transcripts also indicated that transplanted hiPSC microglia upregulated inflammatory cytokine expression, increased phagocytosis, and decreased expression of microglia homeostatic genes and neurotrophic factors (Figure 7. Suppl., Suppl. Table 5). Taken together, these findings further demonstrate that the xenografted human iPSC-derived microglial cells generate functional responses to in vivo injury that closely resembled those of endogenous homeostatic retinal microglia.

**Figure 7.**
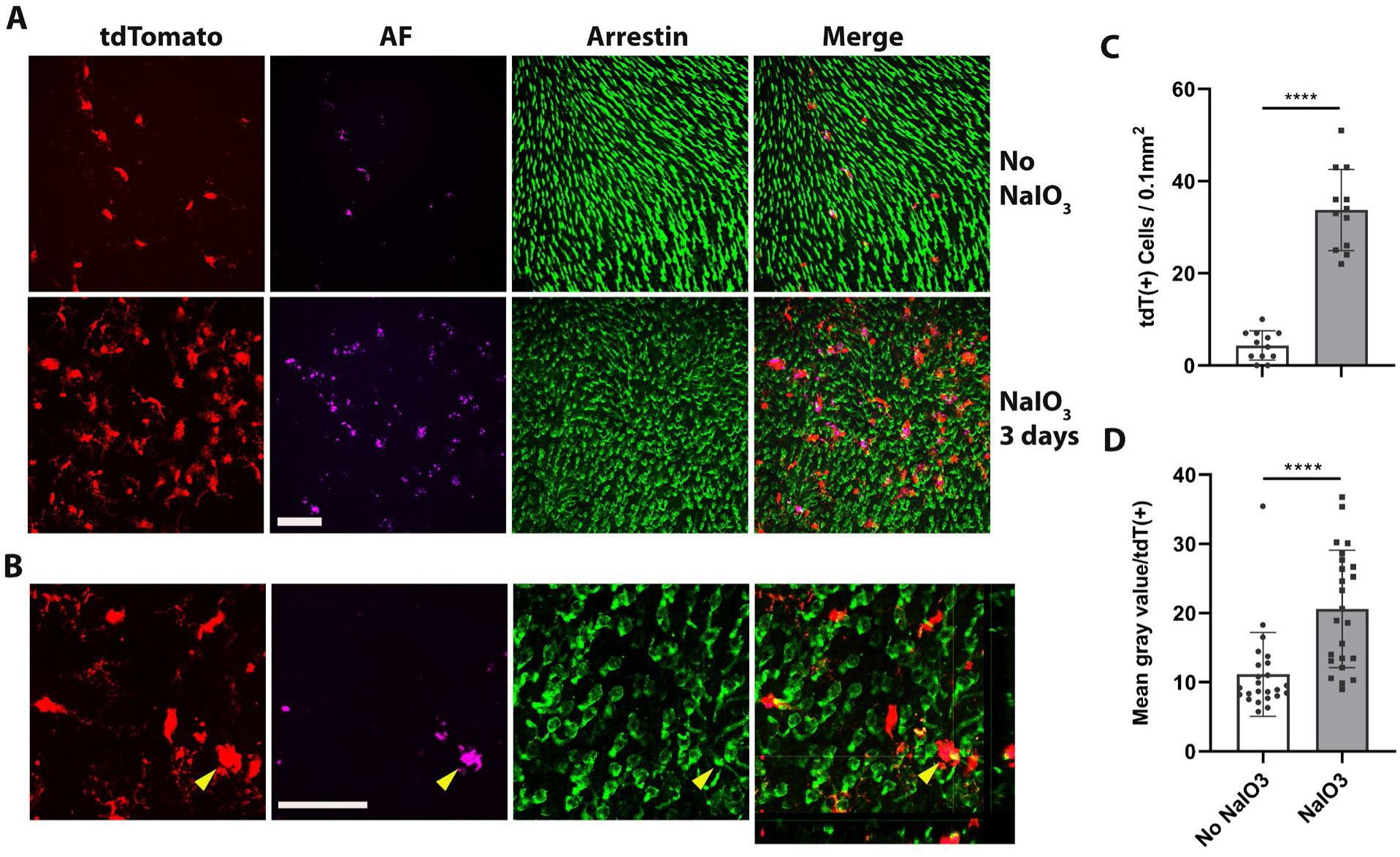
Dyshomeostasis human iPSC-derived microglial cells in the mouse retina phagocytose dead photoreceptor cells/debris after RPE cell injury. (A) Dyshomeostasis human microglial cells (tdtomato+) accumulated in the photoreceptor cell layer after 3 days of NaIO3-induced RPE cell injury compared with no NaIO3 administration. The photoreceptor cells stained with cone arrestin (green), autofluorescent showed in magenta. Scale bar = 60µm. (B) High magnificent and side view image showed human microglial cells (red) co-labeled with photoreceptor cells arrestin staining (green) after 3 days of NaIO3 injury. The yellow triangle showed the colocalized tdT+ human microglia cell and arrestin+ cone photoreceptor cell. Scale bar =40µm. (C) The number of human microglial cells in the photoreceptor layer; (D) The mean gray value of autofluorescence in each human microglia cell. **** P<0.0001.

## Discussion

Microglial cells are instrumental in the development and progression of numerous CNS diseases. For instance, in Alzheimer’s disease, microglia are enriched in over 50% of associated gene loci implicated in AD risk (Hasselmann et al., 2020). Similarly, in Age-related Macular Degeneration, 57% of 368 genes, located close to 52 AMD gene loci, are expressed in retinal microglial cells (Fig. discussion suppl B), and 52% of them are highly expressed in these cells (Fig. discussion suppl A & B; Fritsche et al., 2016; Ma et al., 2013; Den et al., 2022). Understanding microglia cell functions is essential for investigating disease mechanisms and identifying accurate therapeutic targets. Most of our current knowledge about microglial cells comes from rodent studies. However, genetic and functional differences exist between murine and human microglia (Galatro et al., 2017; Gosselin et al., 2017; Smith & Dragunow, 2014). Hence, more in-depth knowledge of human microglial cells in vitro and in vivo is required.

Human-induced pluripotent stem cells (iPSC) offer promising prospects for many retinal research fields (Zhong et al., 2014; Leach et al.,2016; Tanaka et al., 2016). For over a decade, macrophage/microglial cells have been differentiated using human iPSC (Karlsson et al., 2008; Pocock et al., 2018). An abundance of hiPSC-derived microglial cells, with defined genomic background, and easy manipulation of hiPSC offer substantial benefits in various research areas.

Under in vivo physiological conditions, microglial cells exhibit a tile-like arrangement, without overlap. They try to maintain this property in culture, allowing overgrown cells to float out to the medium. Based on this phenomenon and a variety of established microglia cell differentiation protocols (Muffat et al., 2016; Pandya et al., 2017; Abud et al., 2017; Douvaras et al., 2017; Haenseler et al., 2017; Takata et al., 2017), We chose to employ the myeloid progenitor/microglia cell floating culture method (described in Van et al., 2013, and Haenseler et al., 2017) due to its simplicity, efficiency, and consistency. This approach facilitates the generation of a large and uniform population of myeloid progenitor cells, with over 98.6% expressing the CD34+ marker, sustained over a three-month period. These progenitor cells differentiate into pure microglial cells (>98.5% P2ry12+), bearing a profile rich in microglia genes, and demonstrate characteristics similar to native microglia in physiological CNS tissue. A comparison of the signaling pathway to myeloid cells reveals the central hubs of signaling as IL6, IL1b, and the stat3 pathway (Fig.2D), which are key to microglia functioning during inflammation. The differentiation protocol employing floating myeloid progenitor cells produces a significant number of CNS resident-like microglial cells. These hiPSC-derived microgial cells respond robustly to LPS stimulation and demonstrate phagocytic activity, mimicking primary cultured retinal microglial cells. Gene expression profile comparison between hiPSC-microglial cells and fetal/adult brain microglia revealed that hiPSC-derived microglial cells are comparable to fetal and adult microglial cells but are far away from monocytes and inflammatory monocytes (Fig.2. Suppl. B, C and D; Suppl. table 1 and table 2). Therefore, they effectively replicate resident microglia characteristics (Ulland et al., 2018; Wang et al., 2020; Shi et al., 2022; Guo et al., 2022).

CNS disease treatment strategies include gene regulation, gene delivery (Neumann et al., 2006; Beutner et al., 2013), rejuvenation (Karlstetter et al., 2015; Elmore et al., 2018), or replacement of dysfunctional microglial cells (Willis et al., 2020; Xu et al., 2020; Han et al., 2020; Shibuya et al., 2022). hiPSC-derived microglial cells offer a potentially unlimited source for cell replacement therapy. The in vivo functionality of these cells was verified through xenografting into adult Rag2-/-; IL2rg-/-;hCSF1+/+ mice. The grafted microglial cells integrated into the right location of the retina, expressing microglia signature genes and formed new homeostasis with resident microglial cells for eight months. Homeostatic hiPSC-derived microglia responded to RPE cell injury like host retinal microglial cells, marking the success of the xenotransplantation model (Guilliams et al., 2017) and highlighting the potential for hiPSC-derived microglial cells in retinal microglia cell replacement therapy, such as in AMD.

Several exogenous microglia cell replacement techniques have been investigated, with these methods varying based on the type of donor cells used and the age of the recipient. Initial microglia cell replacement studies started from the transplantation of hematopoietic stem cells (HPSC) (Larochelle et al., 2016; Xu et al., 2020; Hohsfield et al., 2020). However, even after differentiation in local tissue, HPSC-derived microglial cells maintained some gene expressions distinct from the original resident microglial cells (Lund et al., 2018), warranting further exploration of these HPSC-derived microglial cells’ unique characteristics. Another method involves the use of newborn mice as recipients, with iPSC or stem cells-derived microglial cells as the donors (Mancuso et al., 2019; Xu et al., 2020). This approach does not require the depletion of resident microglial cells, and the grafted cells can infiltrate the brain tissue, distributing similarly to the original resident microglial cells. While suitable for examining microglia cell function in various backgrounds, the clinical applicability of this method remains limited.

A third transplantation approach involves adult recipients receiving iPSC or stem cell-derived microglial cells after resident microglia cell depletion (Chadarevian et al., 2023). Since this technique requires a microglial-empty niche, resident microglia must be depleted or relocated from the retina, typically done by using CSF1R inhibitors. To prevent these inhibitors from affecting the grafted cells, investigators have tried using modified CSF1R microglial cells as donor cells (Chadarevian et al., 2023). In our research, we found a two-day recovery period with normal food intake sufficient to clear the CSF1R inhibitor, allowing grafted cells to integrate into microglia empty niche. In this study, we used the subretinal xenotransplantation method, though alternative transplantation routes such as intravenous injection, vitreous, and suborbital space delivery warrant further exploration.

In conclusion, hiPSCs can be differentiated into microglial cells through a simplified common pathway and factors, although the more precise differentiation factors still need further investigation under in vitro conditions. For instance, the microglia signature gene, TMEM119, shows low expression in hiPSC-derived microglia single-type cell culture conditions. Interestingly, we discovered that a medium composed of mix cultured embryoid bodies and microglia progenitor cells fosters further differentiation of microglial cells. The humanized mouse model established through xenotransplantation serves as a reliable tool for studying the functions of various hiPSC-derived microglial cells. This model offers a valuable platform for investigating disease mechanisms and evaluating the therapeutic effects of xenografted hiPSC-derived microglial cells from variety of backgrounds.

## EXPERIMENTAL PROCEDURES

### Experimental animals and PLX-5622 treatment

In vivo experiments were conducted according to protocols approved by the Institutional Animal Care and Use Committee (National Eye Institute Animal Care and Use Committee) and adhered to the Association for Research in Vision and Ophthalmology (ARVO) statement on animal use in ophthalmic and vision research. Rag2^-/-^;IL2rg^-/-^;hCSF1^+/+^ transgenic mice were obtained from Jackson Laboratories (Stock #17708). Animals were housed in a National Institutes of Health (NIH) animal facility under a 12-h light/dark cycle and fed standard chow. Genotype analysis by sequencing revealed that the rd8 mutation was absent in the *Crb1* gene. To deplete retinal microglia, two-month old experimental animals were administered a diet containing PLX-5622 (Plexxikon, at 1200 parts per million), a potent and selective inhibitor of the CSF1R previously demonstrated to deplete the majority of microglia in the mouse brain (Dagher et al., 2015) and retina (Zhang et al., 2018). Animals were maintained continuously on the PLX-5622 diet for 10 days and then switched back to standard chow.

### Human iPSC Culture

Five separate human iPSC lines were used in this study. The KYOUDXR0109B (201B7, ATCC:ACS1023) hiPSC line was generated in Yamanaka Lab from fibroblasts isolated from a healthy female donor and reprogrammed by the expression of OCT4, SOX2, KLF4 and MYC using retroviral transduction. The lines of NCRM5-AAVS1-CAG-EGFP (clone 5), ND2-AAVS1-iCAG-tdTomato (clone 1), and NCRM6 and MS19-ES-H were obtained from the NHLBI iPSC Core Facility of National Heart, Lung and Blood Institute (NHLBI). NCRM5-AAVS1-CAG-EGFP is an EGFP-expressing reporter iPSC line with CAG-EGFP targeted mono-allelically at the AAVS1 safe harbor locus in NCRM5 iPSCs. These iPSCs were reprogrammed from CD34+ PBMCs from a healthy male donor. ND2-AAVS1-iCAG-tdTomato is a tdTomato-expressing reporter iPSC line with insulated CAG-tdTomato targeted mono-allelically at the AAVS1 safe harbor locus in ND2 iPSCs. These iPSCs were reprogrammed using healthy male fibroblast cells. Microglia differentiated from these two reporter lines were used for the xenotransplantation experiments. The NCRM6 iPSC line was reprogrammed from CD34+ PBMCs from a healthy female donor with episomal iPSC reprogramming vectors (Thermo Fisher). The MS19-ES-H line was reprogrammed from PBMCs from a healthy female donor with a Cytotune Sendai Virus kit (Thermo Fisher).

The derivation and characterization of iPSC lines were conducted at the ATCC and NHLBI iPSC Core Facility. Cells were cultured on Geltrex-coated (0.2mg/ml, Gibco, #A1413302) 6-well plates using 1x mTeSRTM-1 medium. Passaging was performed following dissociation with TrypLE Express enzymatic digestion (Gibco by Life Technologies). Upon initial plating, cells were cultured in medium containing 3µM Rho-kinase inhibitor (Y-27632, Abcam). The medium was completely refreshed every day. Cells reaching 70% confluence were either passaged or cryopreserved in Stem Cell Freezing Media (mFreSR™, StemCells, Catalog # 05854).

### Myeloid progenitor cell differentiation and microglial cell maturation

The protocol for the differentiation of hiPSC cells to myeloid progenitors and then to microglia were adapted and modified from those described previously by the Cowley laboratory (Van et al., 2013, Haenseler et al., 2017). The first step of the protocol to enable embryoid body (EB) formation employed the Spin-EBs formation method performed in AggreWellsTM800 microwell culture plates (Stemcell Technologies, Catalog # 34825). Briefly, 1 mL of mTeSRTM-1 spin-EB medium was added into each culture well and centrifuged at 3000G for 2 minutes. Subsequently, 4x10^6^ iPSCs in 1ml of medium were added to the well of spin-EB plate and then centrifuged at 800 rpm for 3 minutes. The plate was incubated at 37°C, 5% CO_2_ for four days, then 1 mL medium was replaced in a drop-wise manner every day for the next 4 days with an EB medium: mTeSR1 medium (STEMCELL Technologies) containing 50 ng/mL BMP-4 (GIBCO-PHC9534), 20 ng/mL human stem cell factor (SCF, Miltenyi Biotec), 50 ng/mL vascular endothelial growth factor (hVEGF, GIBCO-PHC9394).

In the second step of myeloid progenitor differentiation, the resulting EBs were harvested with a 40µm filter column. Approximately 150-200 EBs were transferred into a 75-cm^2^ flask containing myeloid cell differentiation medium: TheraPEAK™ X-VIVO™-15 Serum-free Hematopoietic Cell Medium (Lonza, Cat#: BEBP04-744Q) containing 100 ng/mL M-CSF (Invitrogen), 25 ng/mL IL-3 (R&D), 2 mM Glutamax supplement (Invitrogen), 1xN2 supplement (Thermofisher Scientific, Cat#17502048)). Two-thirds of the media volume in the culture flask was replaced every five days for 2-3 weeks.

In the third step of microglia differentiation, the non-adherent floating cell layer consisting of differentiated myeloid progenitor cells was harvested from the supernatant and transferred into the 6-well plate and allowed to settle and adhere overnight. Two-thirds of the medium volume was removed and replaced with a microglia cell differentiation medium: DMEM/F12 medium (Gibco, #11330) containing 50 ng/mL M-CSF (Invitrogen), 100 ng/mL IL-34 (Peprotech), 10 ng/mL TGFb1 and 2-5ng/ml TGFb2 (both from R&D Systems), 20ng/ml CX3CL1 (Peprotech) ,1x N2 supplement and 2 mM Glutamax supplement). The culture was maintained for 2 weeks after which the differentiated iPSC-derived microglia were harvested for analysis or for cell transplantation.

### Phagocytosis Assay

To assess microglia phagocytosis in vitro, the following bioparticles were employed as targets: 1) pHrodo™ Red E. coli BioParticles Conjugate for Phagocytosis (ThermoFisher Scientific, Cat#P35361); 2) pHrodo™ Red Zymosan Bioparticles™ Conjugate for Phagocytosis (ThermoFisher Scientific, Cat#P35364); 3) bovine rod photoreceptor outer segments (POS, Invision Bioresources, Cat#98740). Bovine POS were diluted in serum-free DMEM/F12 (1:1; Gibco) to a concentration of 10^6^ segments/ml and fluorescently labeled with the lipophilic dye DiI (Vybrant Cell-Labeling Solutions; Invitrogen) according to the manufacturer’s instructions. Each phagocytosis assay employed 1x10^5^ POSs and 2mg/ml pHrodo™ Red E. coli membrane/Zymosan BioParticles.

For the assay, harvested floating myeloid cells were transferred into 4-well chamber slides (Thermo Fisher). For phagocytosis assessment of myeloid cells, the cells were cultured for one day and challenged with bioparticles. To assess iPSC-derived microglia, the cells were cultured in a microglia cell differentiation medium for two weeks and then challenged. In the assay, bioparticles were added to the 100µl serum-free DMEM/F12 medium in the slide chamber, incubated for one hour at 37°C, washed 3 times with PBS, and then fixed in 4% paraformaldehyde (PFA) for 20 minutes. Fixed cells were immunostained with antibodies to IBA1 and P2RY12 and counterstained with DAPI. Stained cells were imaged with an Olympus 1000 confocal microscope, and image analysis was conducted using Image J software (NIH).

### mRNA and protein analysis following LPS challenge in vitro

Differentiated hiPSC-derived microglia cultured in in 6-well plates were stimulated with LPS at 100 ng/mL for 6 and 24 hr. Microglia stimulated for 6 hrs were collected in RNAlater solution (Thermo Fisher) and stored at -80°C for further QRT-PCR analysis. Microglia stimulated for 24 hrs were collected for protein quantification. After the medium was collected, the cells in the well were washed with 1XPBS, and then 200µl of RIPA lysis buffer with proteinase inhibitor cocktail (Calbiochem) was added; the cells were removed by scraping, collected into 1.5ml Eppendorf tube, and then homogenized with sonication (Sonicator 125 Watts, Qsonica) at 4°C. After sonication and centrifugation, total protein concentration was measured (BCA protein assay kit; Pierce). Levels of individual cytokines were determined using a Milliplex bead assay kit (Milliplex MAP human cytokine/chemokine magnetic bead panel, #MCYTOMAG-70K; Millipore) and involving the Luminex MAPIX system with data analysis using xPONENT 4.2 software (Luminex). The cytokines analyzed included *IL1α, IL1β, IL6, IL8, TNFα, CXCL10, CCL2, CCL3, CCL4, IL10*.

### mRNA expression analysis by quantitative RT-PCR

mRNA expression was quantitated using quantitative reverse transcription-PCR (qRT-PCR). Harvested cells were lysed by trituration and homogenized using QIAshredder spin columns (Qiagen). Total RNA was isolated using the RNeasy Mini kit (Qiagen) according to the manufacturer’s specifications. First-strand cDNA synthesis from mRNA was performed using qScript cDNA SuperMix (Quanta Biosciences) using oligo-dT as primer. qRT-PCR was performed using an SYBR green RT-PCR kit (Affymetrix), using the Bio-Rad CFX96 Touch™ Real-Time PCR Detection System under the following conditions: denaturation at 95 °C for 5 min, followed by 40 cycles of 95 °C for 10 s, and then 60 °C for 45 s. Threshold cycle (CT) values were calculated and expressed as fold-induction determined using the comparative CT (2ΔΔCT) method. Ribosomal protein S13 (RPS13) and GAPDH were used as internal controls. Oligonucleotide primers are provided in Suppl. Table 4.

### Transplantation of hiPSC-derived microglia by subretinal injection

Differentiated hiPSC-derived microglia grown in flasks were first washed in 1XPBS before being removed by scraping and collected into a 50ml tube in 5ml PBS. Cell numbers were counted using a cell counter (Countess 3, ThermoFisher). Microglia were collected by centrifugation (5 minutes at 4°C, 200G) and the resulting cell pellet resuspended in 1XPBS at a concentration of 5000 cells/µl for in vivo transplantation via subretinal injection. Experimental animals were given general anesthesia (ketamine 90 mg/kg and xylazine 8 mg/kg) and additional topical anesthesia (0.5% Proparacaine HCL, Sandoz) applied to the injected eye. For the injection, the temporal sclera was exposed by a conjunctival cut-down and a scleral incision made 0.5mm behind the limbus using 32G needle to access the subretinal space. The tip of a blunt 32G needle attached to a Hamilton micro-syringe was introduced through the incision at an angle 5-degrees tangent to the globe and advanced 0.5-1 mm into the subretinal space under a dissecting microscope. Microglial cells (5000 cells in 1µl PBS) were slowly injected from the micro-syringe into the subretinal space using an aseptic technique. Post-procedure, treated eyes were carefully inspected for signs of bleeding or distention and intraocular pressure was monitored using a tonometer (iCare TONOLAB, Finland). In the event that intraocular pressure remained elevated (>20mmHg) and/or the globe appeared distended, a vitreous tap was performed using a 33-G needle to reduce intraocular pressure. In the unlikely event that excessive bleeding is observed, the animal will be examined by a veterinarian or euthanized immediately.

### Immunohistochemical analyses

For immunohistochemical analysis of microglia in vitro, microglia were differentiated in 4-well chambered slides, fixed in 4% PFA for 20 minutes and processed for immunostaining. For in vivo analyses, recipient animals were euthanized by CO_2_ inhalation, and the eyes were enucleated. Enucleated eyes were dissected to form posterior segment eyecups and fixed in 4% paraformaldehyde in phosphate buffer (PB) for 2h at 4°C. Eyecups were either cryosectioned (Leica CM3050S) or further dissected to form retinal flat mounts. Flat-mounted retinas were blocked for 1h in a blocking buffer containing 10% normal donkey serum and 1% Triton X-100 in PBS at room temperature. Primary antibodies included IBA1 (1:500, Wako, #019–19741), anti-mouse Tmem119 (1:500, Synaptic Systems, #400 004), anti-human TMEM119 (1:100, Sigma, #HPA051870), anti-mouse Cd68 (1:200, BioRad, #MCA1957), anti-human CD68 (1:100, R&D, #MAB20401), anti-mouse Cd45 (1:100, Bio-Rad, #MCA1388), anti-human CD45 (1:100, R&D, #FAB1430R), cone arrestin (1:200, Millipore, #AB15282), Ki67 (1:30, eBioscience, #50-5698-82), anti-P2RY12 (1:100, ThermoFisher, #PA5-77671 and Sigma, #HPA014518), CD34 (1:50, eBioscience, #14-0341), hCD11b (1:100, R&D, #FAB1699R), mCD11b (BioRad, Cat#: MCA711G, 1:100), CX3CR1 (1:100, Invitrogen, #61-6099-42), hHLA (Invitrogen, #11-9983-42), PU.1 (Invitrogen, #MA5-15064), TREM2 (1:100, Invitrogen, #702886), glutamine synthetase (1:200, Millipore, #MAB302), PKCa (1:200, Sigma-Aldrich, #p4334), GFAP (1:200, Invitrogen, #13–0300), RBPMS (1:100, Phosphosolutions, 1832-RBPMS), Calbindin (1:5000, Swant, CB-38a), and anti-RFP(1:200, RockLand, 600-401-379-RTU). Primary antibodies were diluted in blocking buffer and incubated at 4°C overnight for retinal sections and at room temperature overnight on a shaker for retinal flat mounts. Experiments in which primary antibodies were omitted served as negative controls. After washing in 1xPBST (0.2% Tween-20 in PBS), retinal samples were incubated for 2 hrs at room temperature with secondary antibodies (AlexaFluor 488-, 568- or 647-conjugated anti-rabbit, mouse, rat, goat and guinea pig IgG) and DAPI (1:500; Sigma-Aldrich) to label cell nuclei. Isolectin B4 (IB4), conjugated to AlexaFluor 568/647 (1:100, Life Technologies), was used to label activated microglia and retinal vessels. Stained retinal samples were imaged with confocal microscopy (Olympus FluoView 1000, or Zeiss LSM 880, or Nikon A1R). For analysis at high magnification, multiplane z-series were collected using 20 or 40 objectives; Confocal image stacks were viewed and analyzed with FV100 Viewer Software, NIS-Element Analysis and Image J (NIH).

### RNAseq analysis

For whole transcriptome analysis, cultured myeloid progenitor cells (MPC) and differentiated microglial cells with and without 0.1µg/ml LPS treatment were harvested in the flasks and 6-well plates, respectively. After harvesting, all samples stored frozen in RNAlater (Roche) solution before RNA extraction using the Qiagen RNA Mini Kit. RNA quality and quantity were evaluated using Bioanalyzer 2100 with the RNA 6000 Nano Kit (Agilent Technologies). The preparation of RNA library and transcriptome sequencing was performed using an external vendor (Novogene, Sacramento, CA). Genes with adjusted p-value <0.05 and log2FC (Fold Change) > 1 were considered differentially expressed. Ingenuity Pathway Analysis (IPA, Qiagen) was employed for canonical pathway and graphical pathway analysis. The microglia gene list was constructed from our previous microarray data from retinal microglial cells and published data (Ma et al. 2013; Bennett et al., 2016 ). The heat map, volcano, and histogram plot were performed using Prism 9.5.1 (Graph-Pad).

### Sodium iodate (NaIO_3_)-induced model of RPE cell injury

Recipient mice 8 months following xenotransplantation were were administered a single dose of NaIO_3_ (Honeywell Research Chemicals) at a dose of 30 mg/kg body weight via intraperitoneal injection. Animals were euthanized 3 and 7 days after NaIO3 injection, and their retinas harvested and subjected to histological and molecular analysis.

### Statistics and reproducibility

All data in the graphical panel represent mean ± standard error. When only two independent groups were compared, significance was determined by a two-tailed unpaired t-test with Welch’s correction. When three or more groups were compared, one-way ANOVA with the Bonferroni post hoc test or two-way ANOVA was used. A P value <0.05 was considered significant. The analyses were done in GraphPad Prism v.5. All experiments were independently performed at least three experimental replicates to confirm consistency in observations across replicates.

## Supporting information

Responses to reviewers

## SUPPLEMENTAL INFORMATION

Supplemental Information includes supplemental figures and legends and supplemental tables.

## AUTHOR CONTRIBUTIONS

WM performed and analyzed data and wrote the manuscript; WT and WL designed and supervised the studies and edited the manuscript. LZ and BX performed in vivo procedures, RNF assisted with imaging procedures and collection, TMR supervised and supported the study, and JZ provided the hiPSC lines and iPSC quality control.

## ACKNOWLEDGMENTS

This study was supported by the National Eye Institute Intramural Research program.

---

**Fig 1. Suppl.**
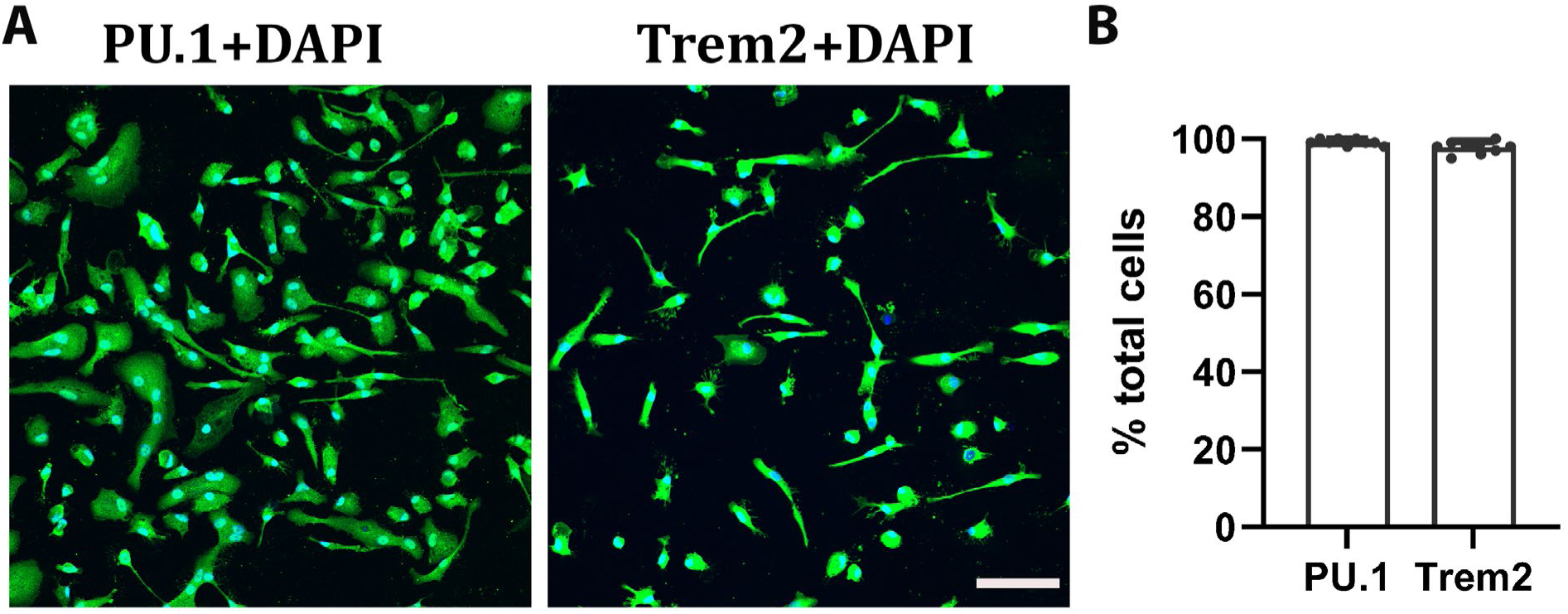
Immunocytochemistry staining with human PU.1 and TREM2(A). Scale bar = 100µm. (B) PU.1 and TREM2 positive cells are 99.7% and 99.4%, respectively, in entire DAPI+ cell counts.

**Fig 2. Suppl A.**
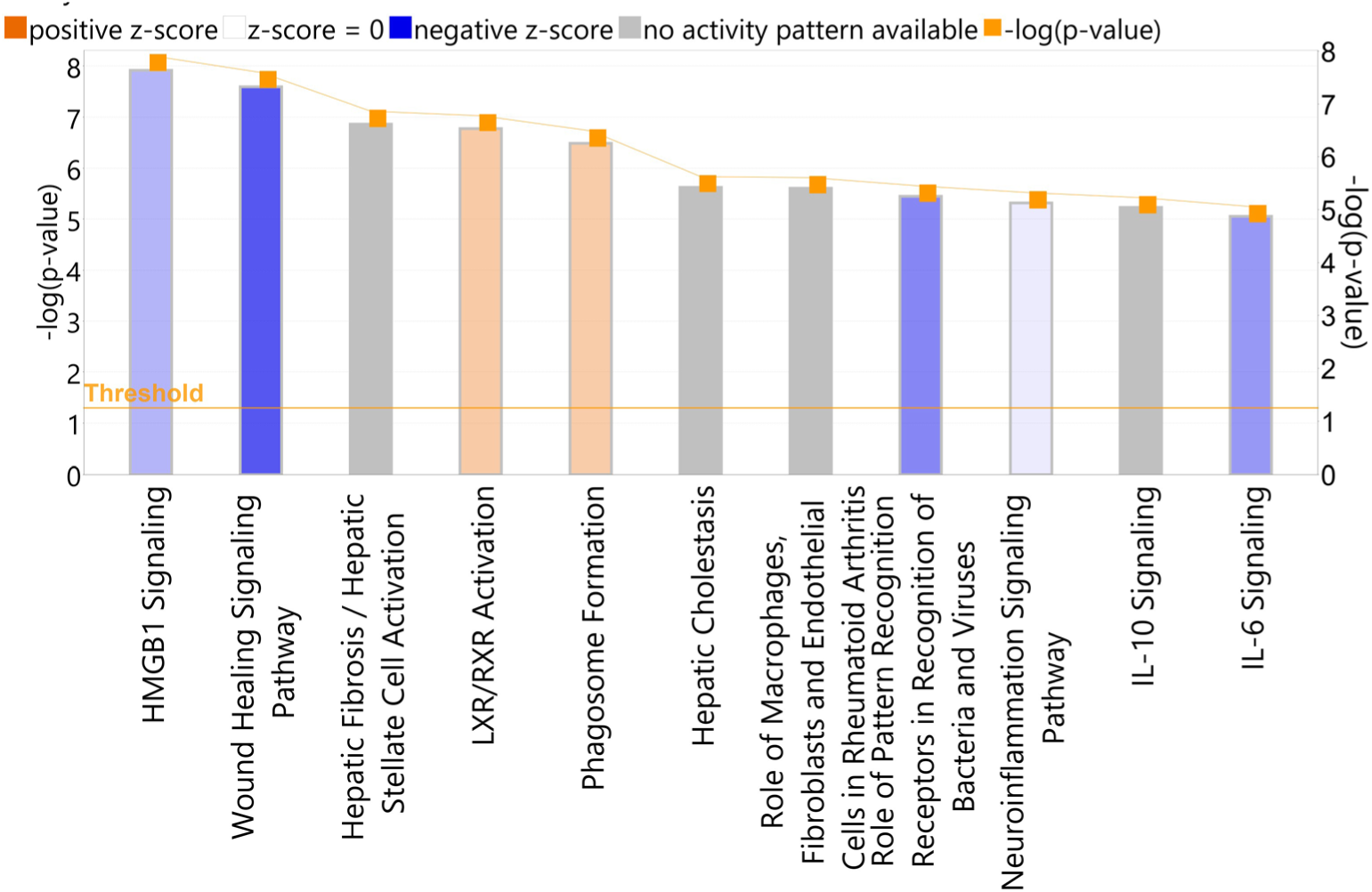
Analysis results of the canonical pathway of completed differentiated hipsc-derived microglia vs. myeloid progenitor cells.

**Fig 2. Suppl B.**
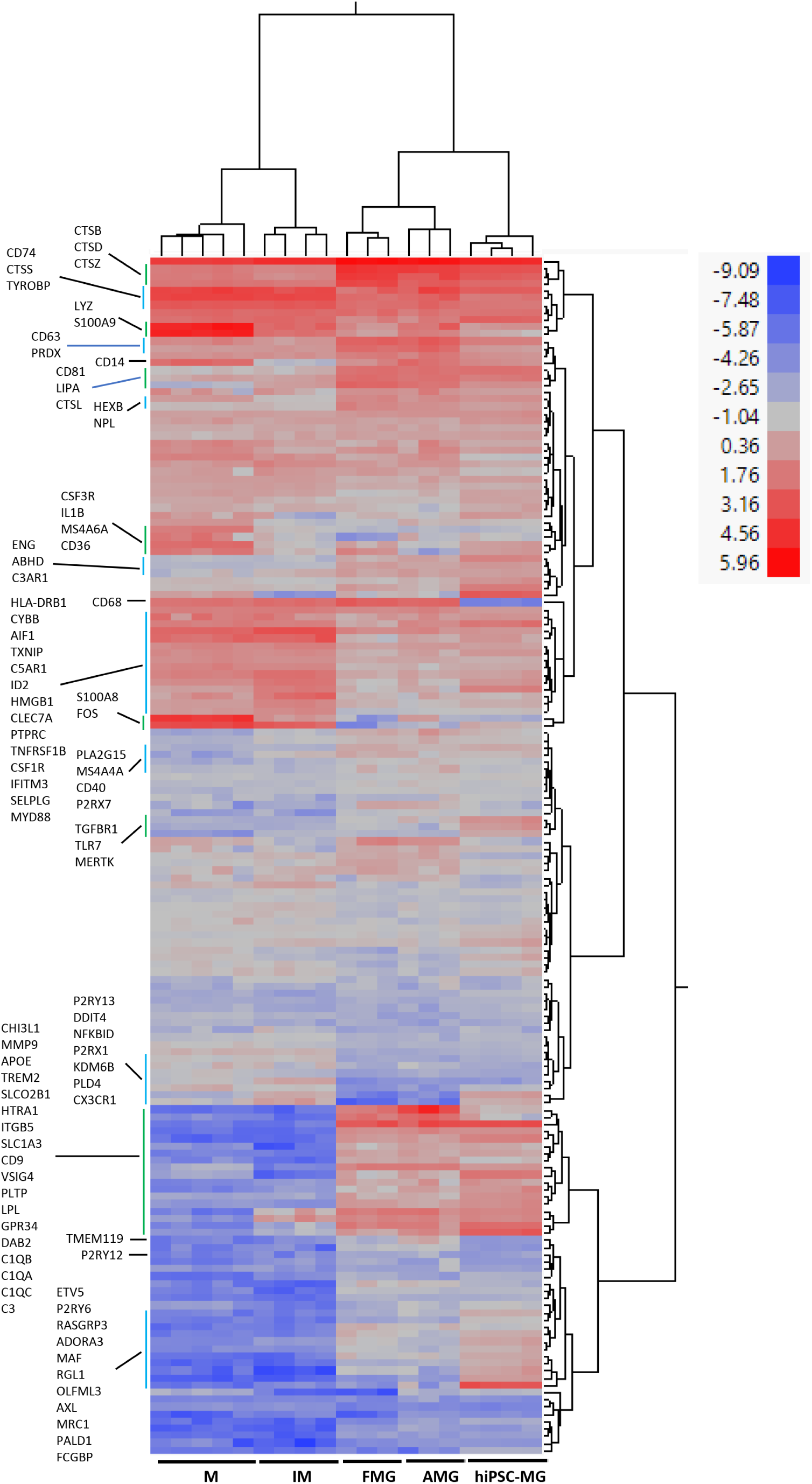
Hierarchical cluster analysis on microglia enriche genes among hiPSC-derived microglial cells (hiPSC-MG) and human adult brain microglia cells (AMG), fetal brain microglia cells (FMG), inflammatory monocytes (IM), monocytes(M). The human microglia gene panel combined our mouse microglia-enriched genes and human microglia-enriched genes (Abud et al., 2017; Muffat et al., 2016; Douvaras et al., 2017; Böttcher et al., 2019; Van et al., 2019). The total microglia enriched gene list contains 203 genes (Suppl. Table 1), which used to be extracted from the gene profile of hiPSC-MG and downloaded human adult microglia(AMG), fetal microglia (FMG), inflammatory monocyte (IM) and monocytes (M) (GSE 178846, Abud et al., 2017). 188 genes (Suppl. table 1) were obtained from both gene profiles. All the gene counts were normalized with 4 human cell housekeeping genes C1orf43, RAB7A, REEP5 and VCP (Eisenberg et al., 2013). The hierarchical cluster was analyzed by JMP (JMP Statistical Discovery LLC.). Results showed that hiPSC-MG is more comparable with AMG and FMG.

**Fig 2. Suppl C.**
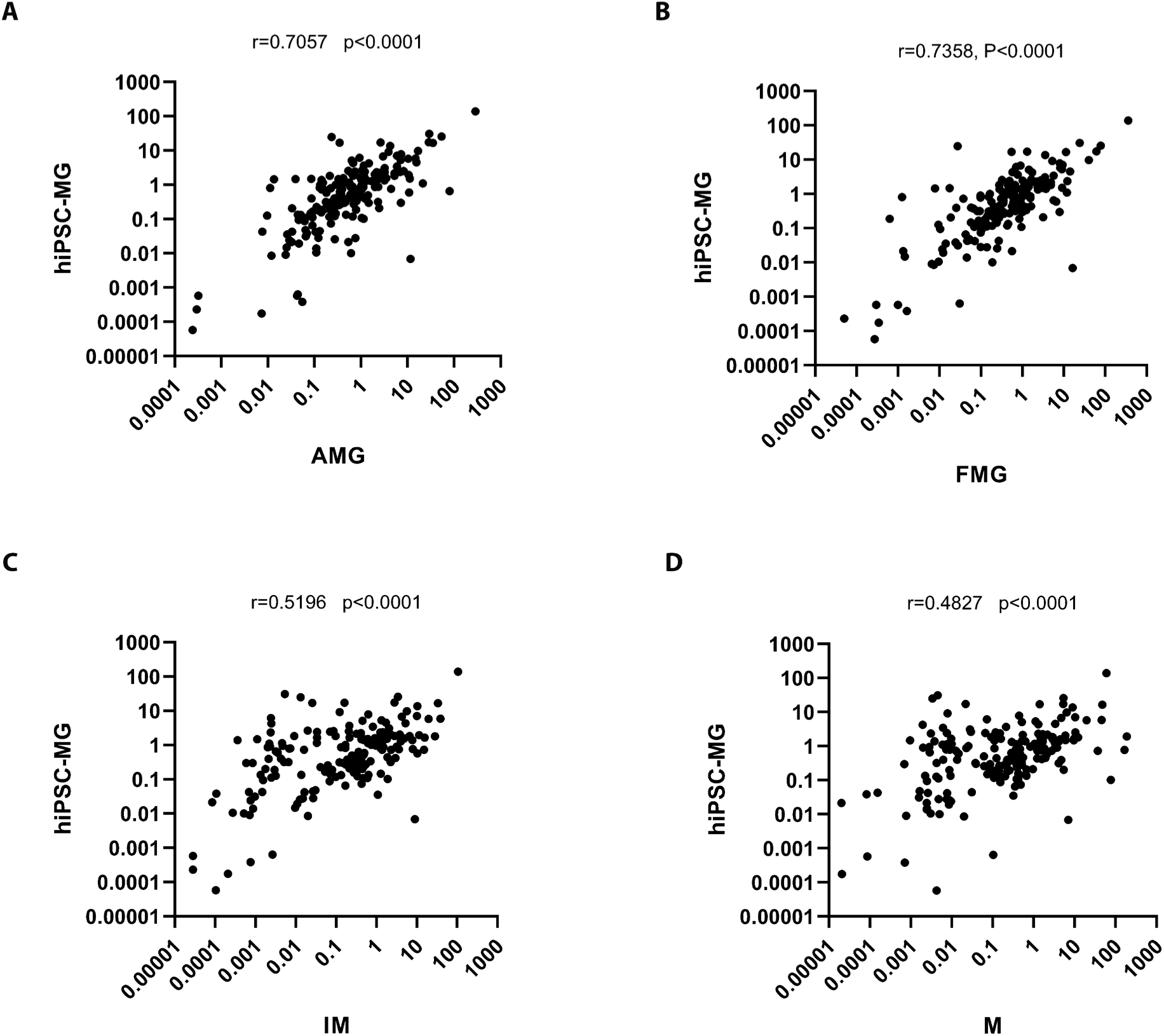
The correlation analysis of human microglia enriched genes among hiPSC-derived microglia cells (hiPSC-MG), human adult brain microglia cells (A), fetal brain microglia cells (B), inflammatory monocytes (C), monocytes (D) respectively. The images and the analysis results (Prism, GraphPad) showed hiPSC-MG correlated significantly with FMG (r=0.7358, p<0.0001) and AMG (r=0.7057, p<0.0001).

**Fig 2. Suppl D.**
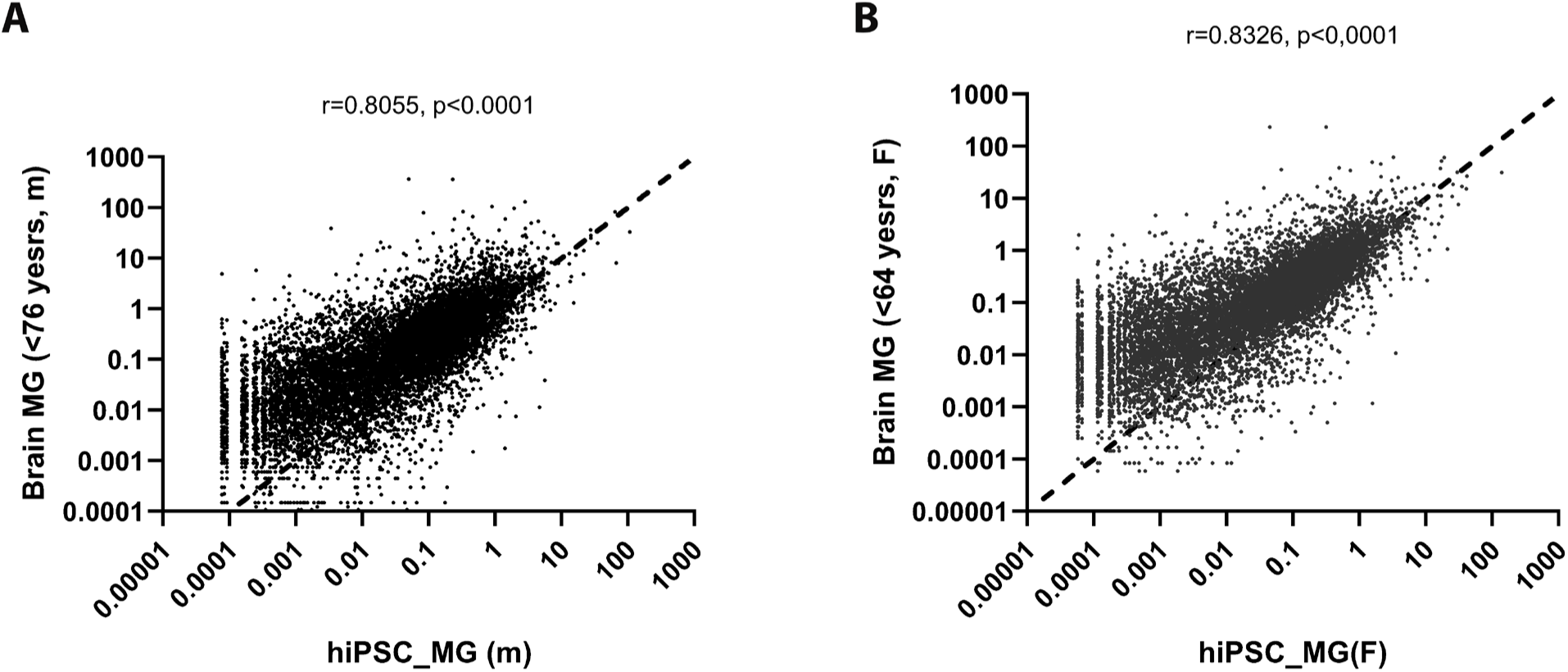
Correlation comparison of entire gene profiles between male (A)/female (B) hiPSC-derived microglial cells and male (A)/female (B) human brain microglia cells (GSE 111972, van der Poel et al. 2019; Suppl. table 2) respectively. The results showed they are significantly correlative (r=0.8055(M), r=0.8326(F), P<0.0001).

**Figure 3 suppl.**
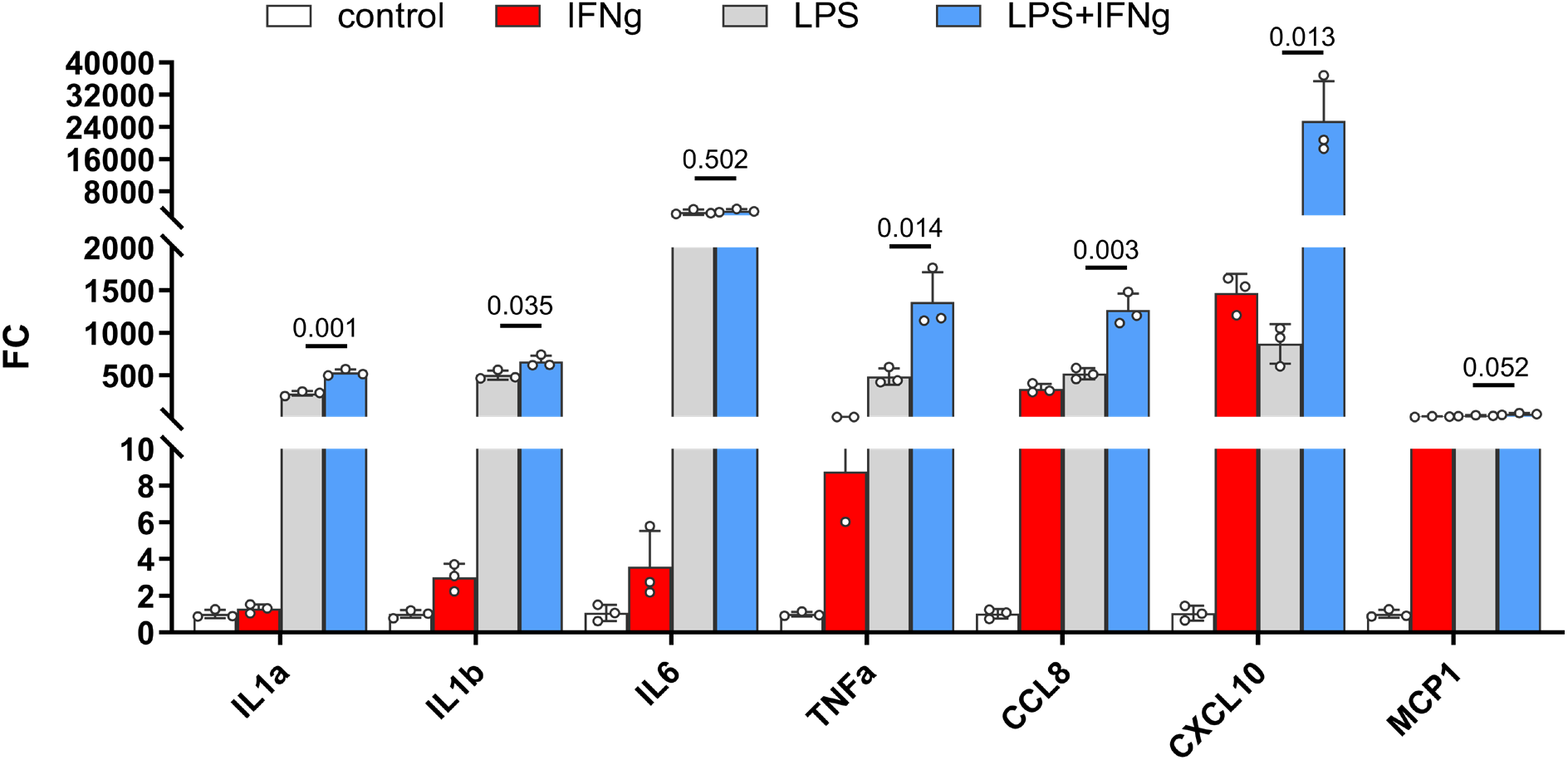
Inflammatory cytokines were produced synthetically by IFNγ and LPS in hiPSC-derived microglial cells. hiPSC-derived microglia cells were cultured in 6-well plates for 14 days, 20ng/ml of human IFNγ and 0.5ug/ml of LPS were added to the wells, respectively, or 20ng/ml of human IFNγ and 0.5ug/ml of LPS were added to the wells together. After 6 hours of incubation, the cells were washed and harvested into a 1.5ml Eppendorf tube for RNA isolation with a NucleoSpin RNA/Protein Mini kit (Macherey-Nagel, #740933.50). The cDNA was synthesized with PrimeScript™ 1st strand cDNA Synthesis Kit (Takara, #6110A). qRT-PCR was performed using a SYBR green RT-PCR kit (Affymetrix), using the Bio-Rad CFX96 Touch™ Real-Time PCR Detection System under the following conditions: denaturation at 95 °C for 5 min, followed by 40 cycles of 95 °C for 10 s, and then 60 °C for 45 s. Threshold cycle (CT) values were calculated and expressed as fold-induction determined using the comparative CT (2ΔΔCT) method. Ribosomal protein S13 (RPS13) and GAPDH were used as internal controls. Oligonucleotide primers are provided in Suppl. Table 4.

**Fig 4. Suppl A.**
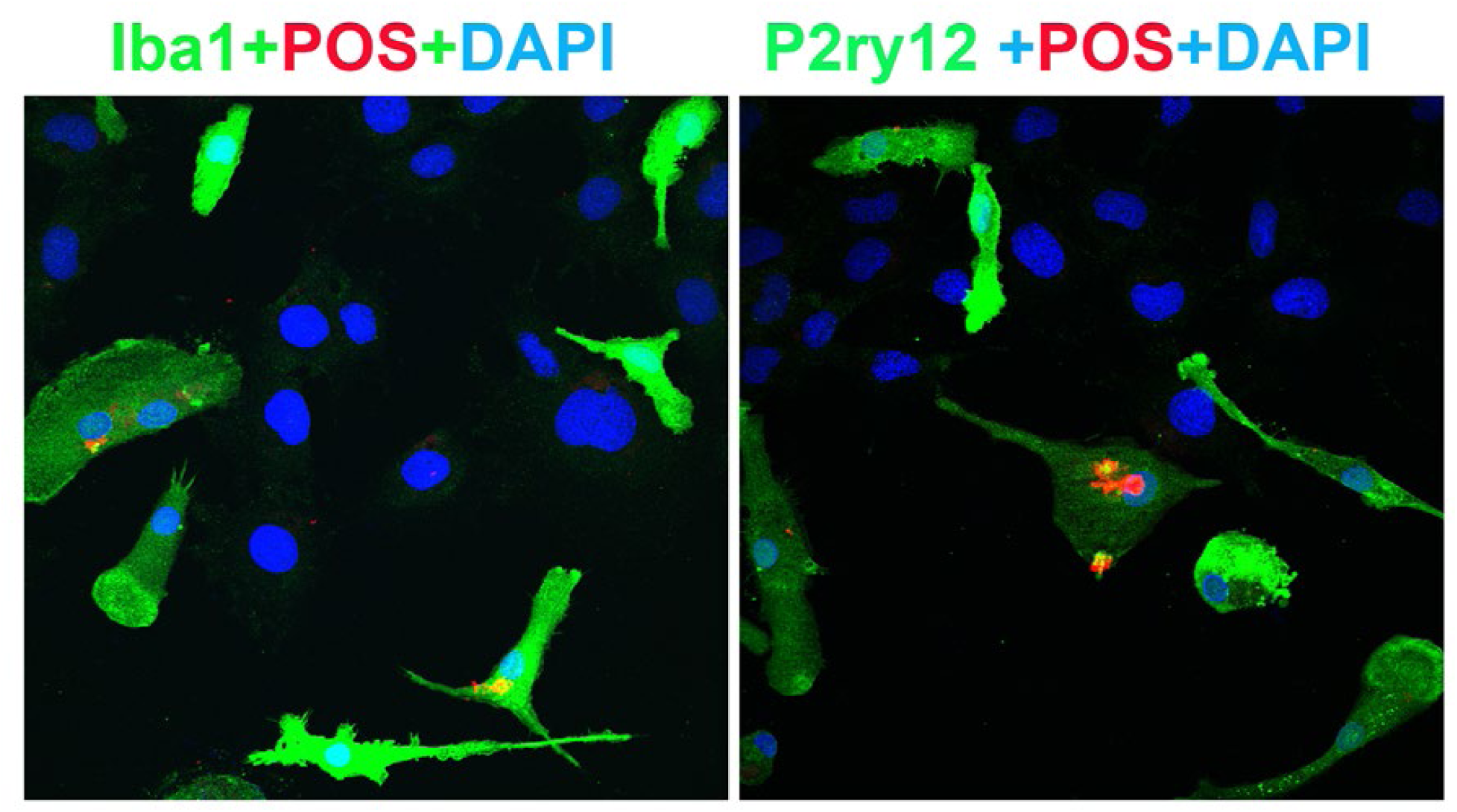
Floating myeloid progenitor cells were cultured in a 2-well slide chamber overnight, and then the DiI-labeled bovine photoreceptor outer segments were added to the chamber and incubated for 1 hr. The cells were fixed with 4% PFA for 20 minutes and stained with Iba1 and hP2ry12, respectively. The DiI-labeled bovine photoreceptor outer segments can only be engulfed by Iba1 or hP2ry12-positive microglia cells. Scale bar = 16µm.

**Figure 4. Suppl B.**
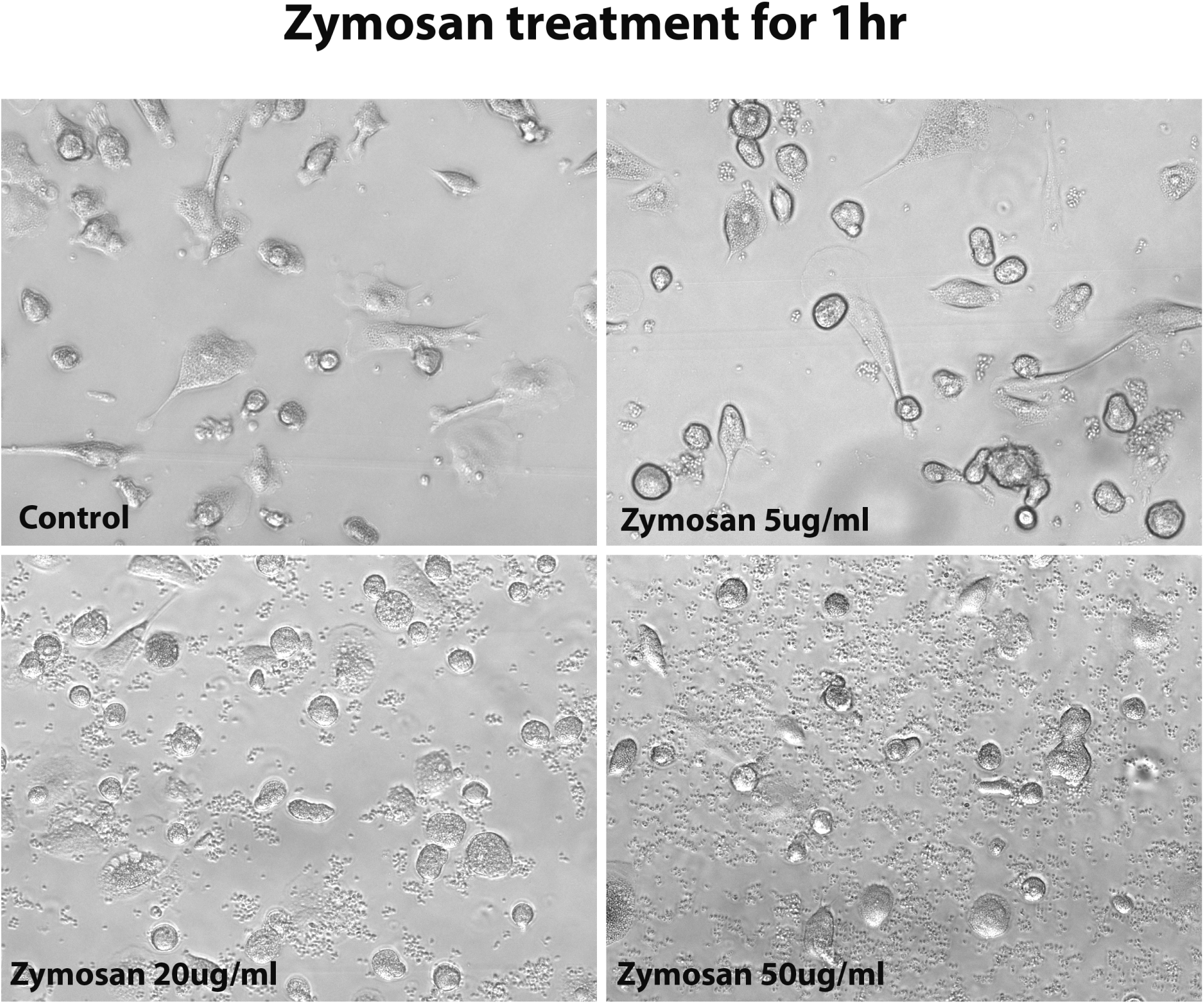
The morphology of hiPSC-derived microglia cells under 1-hour zymosan treatment with different concentrations. After 1 hour of treatment with zymosan, 5ug/ml concentration didn’t change the microglia cell morphology, but over 20ug/ml concentration changed the microglia morphology to an ameboid round shape.

**Fig. 5. Suppl A.**
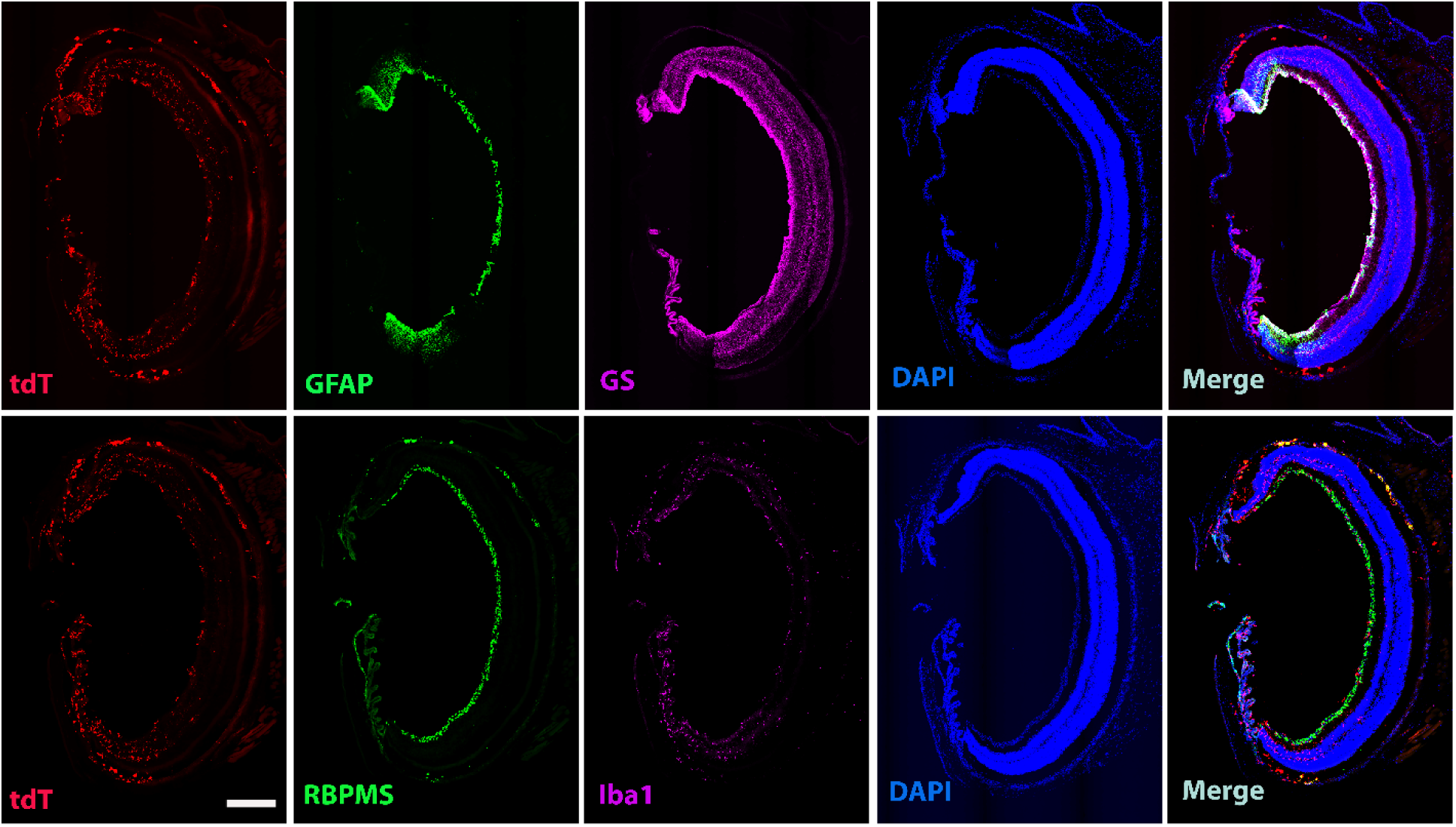
Homeostatic hiPSC-derived microglial cells in the mouse retina do not affect local retinal cells. Entire section images showed GFAP, GS, RBPMS, and Iba1 staining for astrocytes, Müller cells, ganglion cells, and microglia cells in the retina after four months of xenotransplantation. Scale bar = 300µm.

**Fig. 5. Suppl B.**
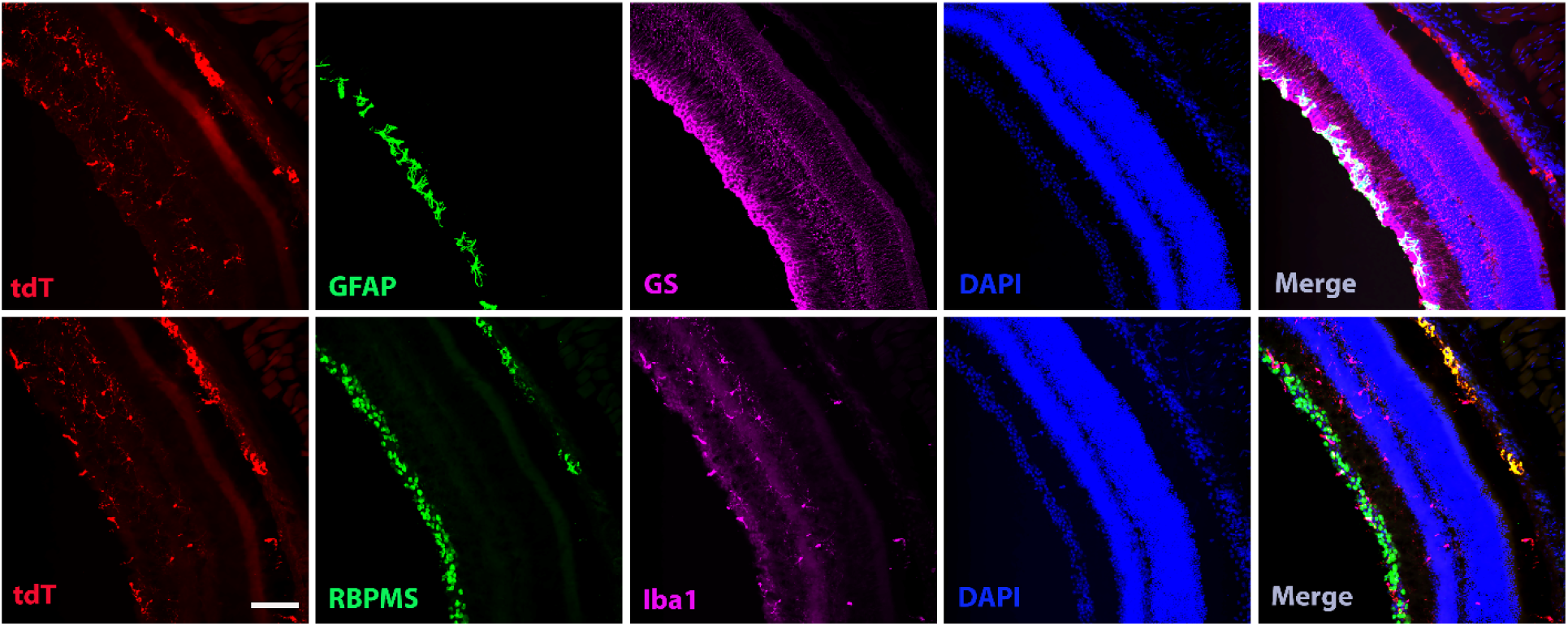
Homeostatic human iPSC-derived microglia cells in the mouse retina do not affect local retinal cells. The partial section high magnification images showed GFAP, GS, RBPMS, and Iba1 staining for astrocytes, Müller cells, ganglion cells, and microglia cells in the retina after four months of transplantation. Scale bar = 100µm.

**Fig. 5. Suppl C.**
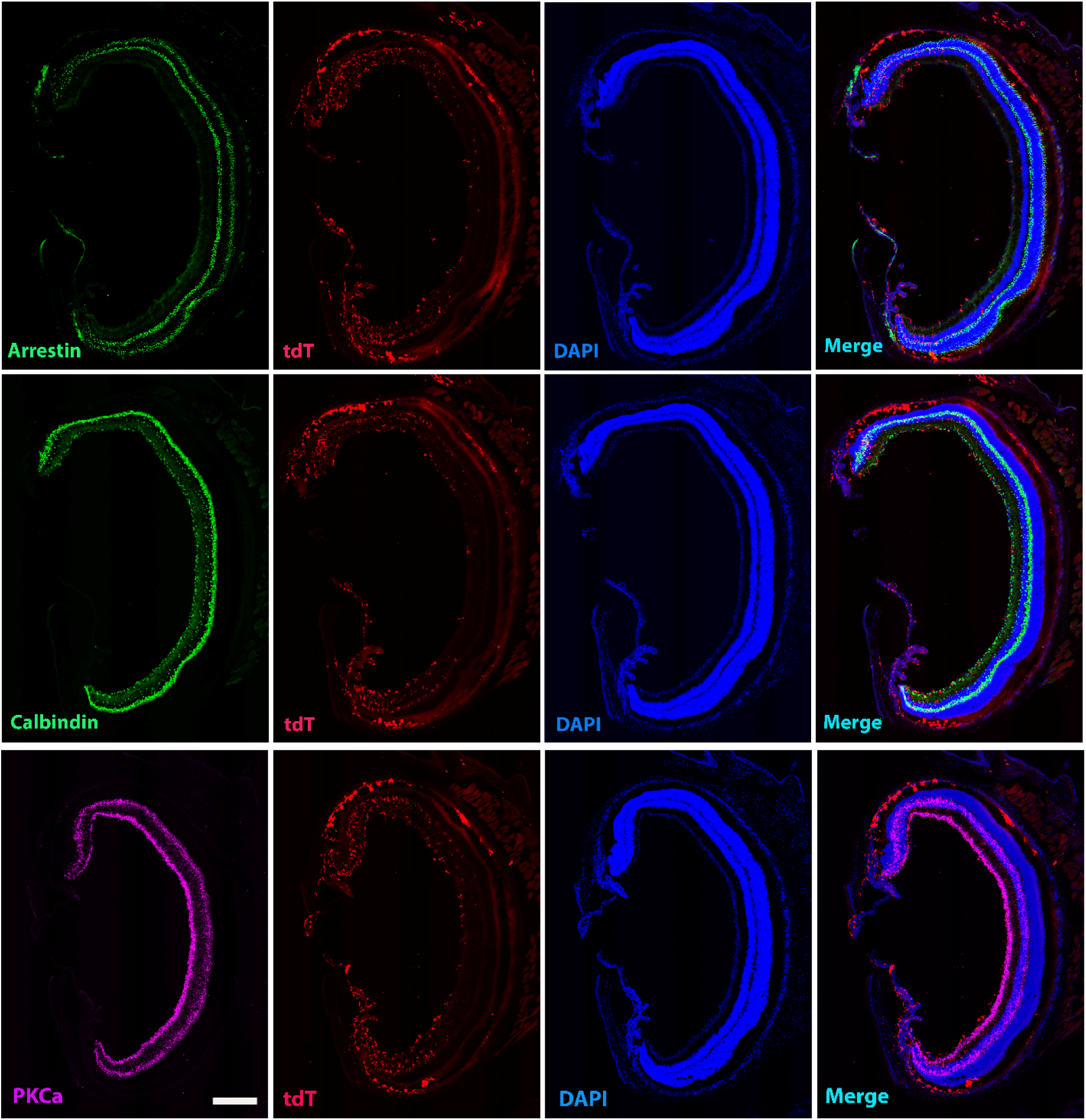
Homeostatic human iPSC-derived microglia cells in the mouse retina do not affect local retinal cells. Entire section images showed arrestin, calbindin, and PKCα staining for cone photoreceptors, horizontal and some amacrine cells, and bipolar cells in the retina after four months of xenotransplantation. Scale bar = 300µm.

**Fig. 5. Suppl D.**
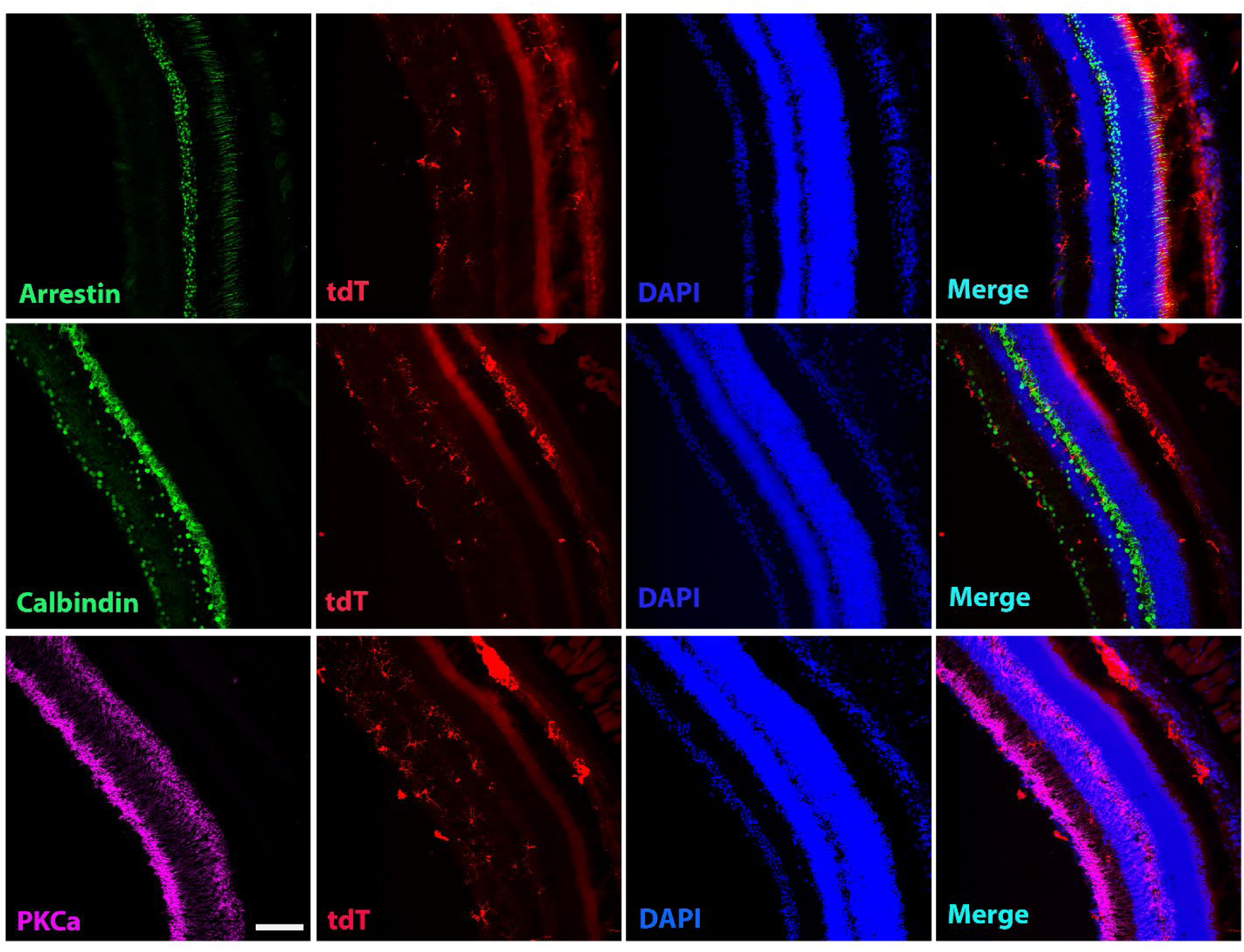
Homeostatic human iPSC-derived microglia cells in the mouse retina do not affect local retinal cells. The partial section of high magnification images showed arrestin, calbindin, and PKCα staining for cone photoreceptors, horizontal and some amacrine cells, and bipolar cells in the retina after four months of xenotransplantation. Scale bar = 100µm.

**Fig. 5. Suppl E.**
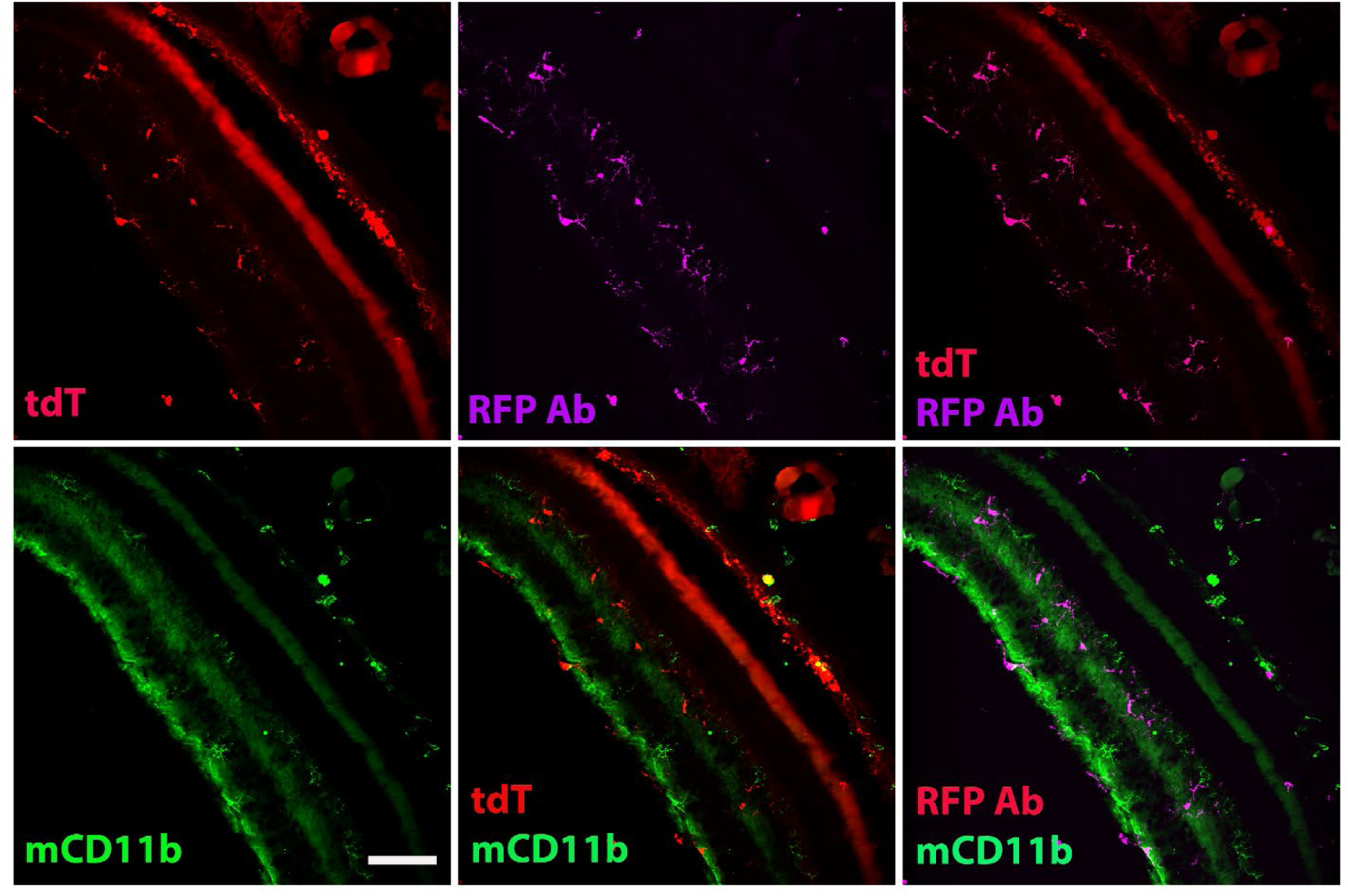
Homeostatic human iPSC-derived microglia cells in the mouse retina do not take over local retinal microglia cells. The section images showed RFP and mouse CD11b staining to determine the tdT+ human microglia cells and local mouse microglia cells in the retina after four months of xenotransplantation. The results showed that the tdT+ human cells colocalized with RFP staining (Far red) but not with mouse CD11b (green) staining. Scale bar = 100µm.

**Fig. 5. Suppl F.**
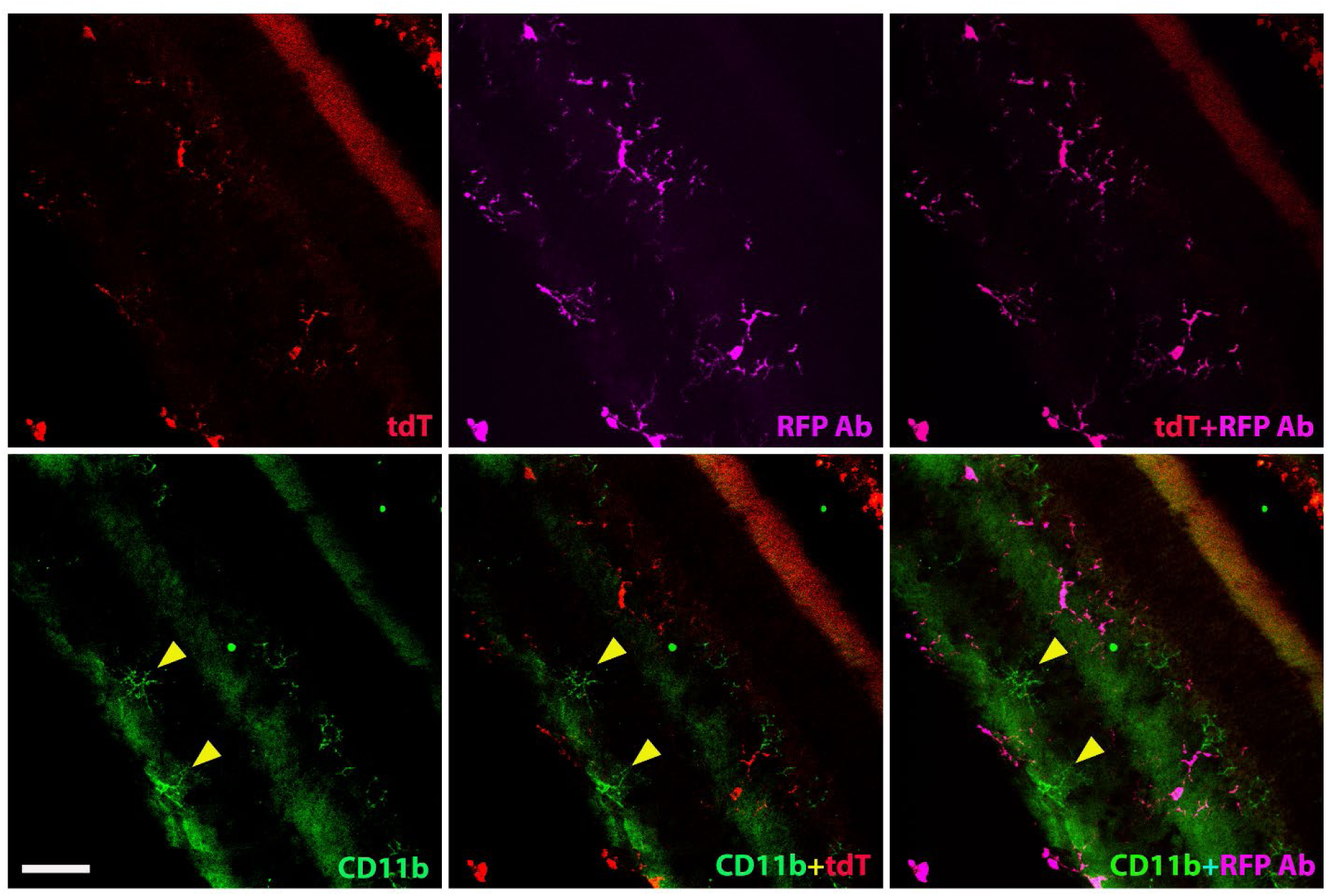
Homeostatic human iPSC-derived microglia cells in the mouse retina do not take over local retinal microglia cells. The magnification images showed RFP and mouse CD11b staining in the retina after four months of xenotransplantation. The results showed that the tdT+ human cells colocalized with RFP staining (Far red) but not with mouse CD11b(green) staining. The triangle marker indicated that the local mouse microglia cells were only stained with mouse CD11b but not colocalized with tdT+ human microglia cells. Scale bar = 50µm.

**Fig. 6. Suppl A.**
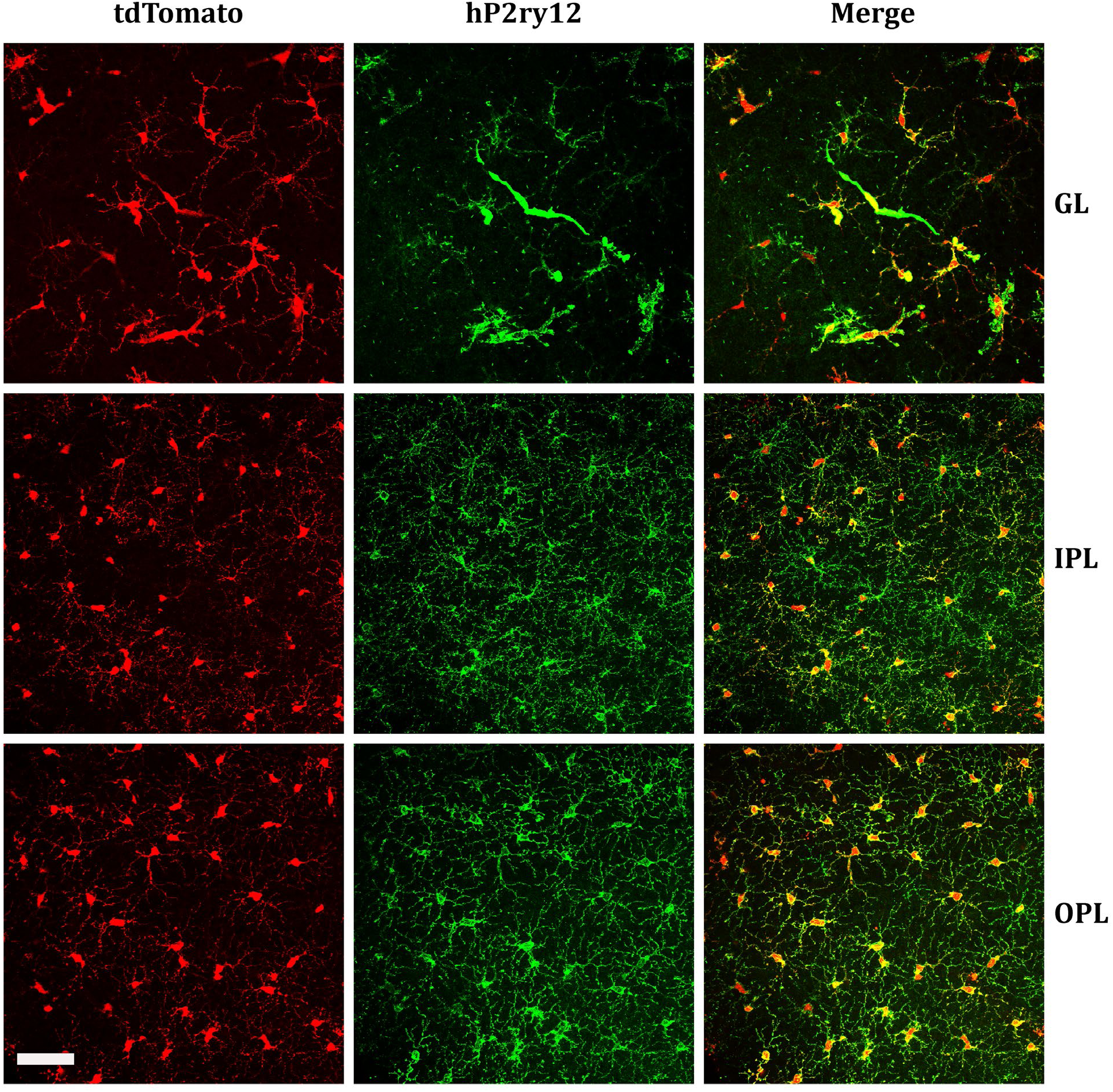
hP2rY12 staining on the retina after eight months of xenotransplantation. The images showed the tdtomato+ human microglia cells colocalized with hP2rY12 staining in GL, IPL, and OPL in the mouse retina. Scale bar = 300µm.

**Fig. 6. Suppl B.**
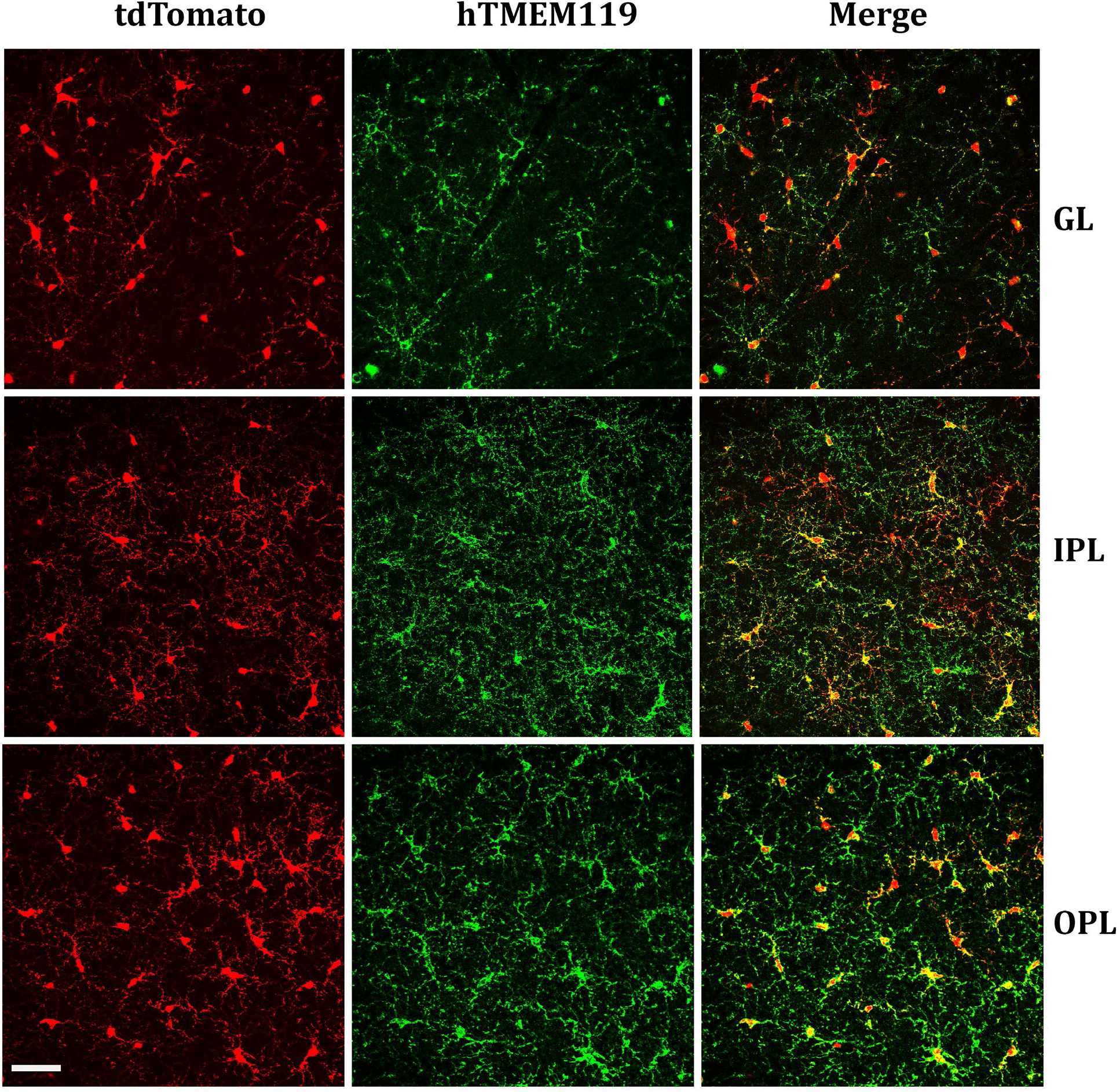
High magnification of hTMEM119 staining on the retina after eight months of xenotransplantation. The images showed the tdtomato+ human microglia cells colocalized with hTMEM119 staining in GL, IPL, and OPL in the mouse retina. Scale bar = 300µm.

**Figure 7. Suppl.**
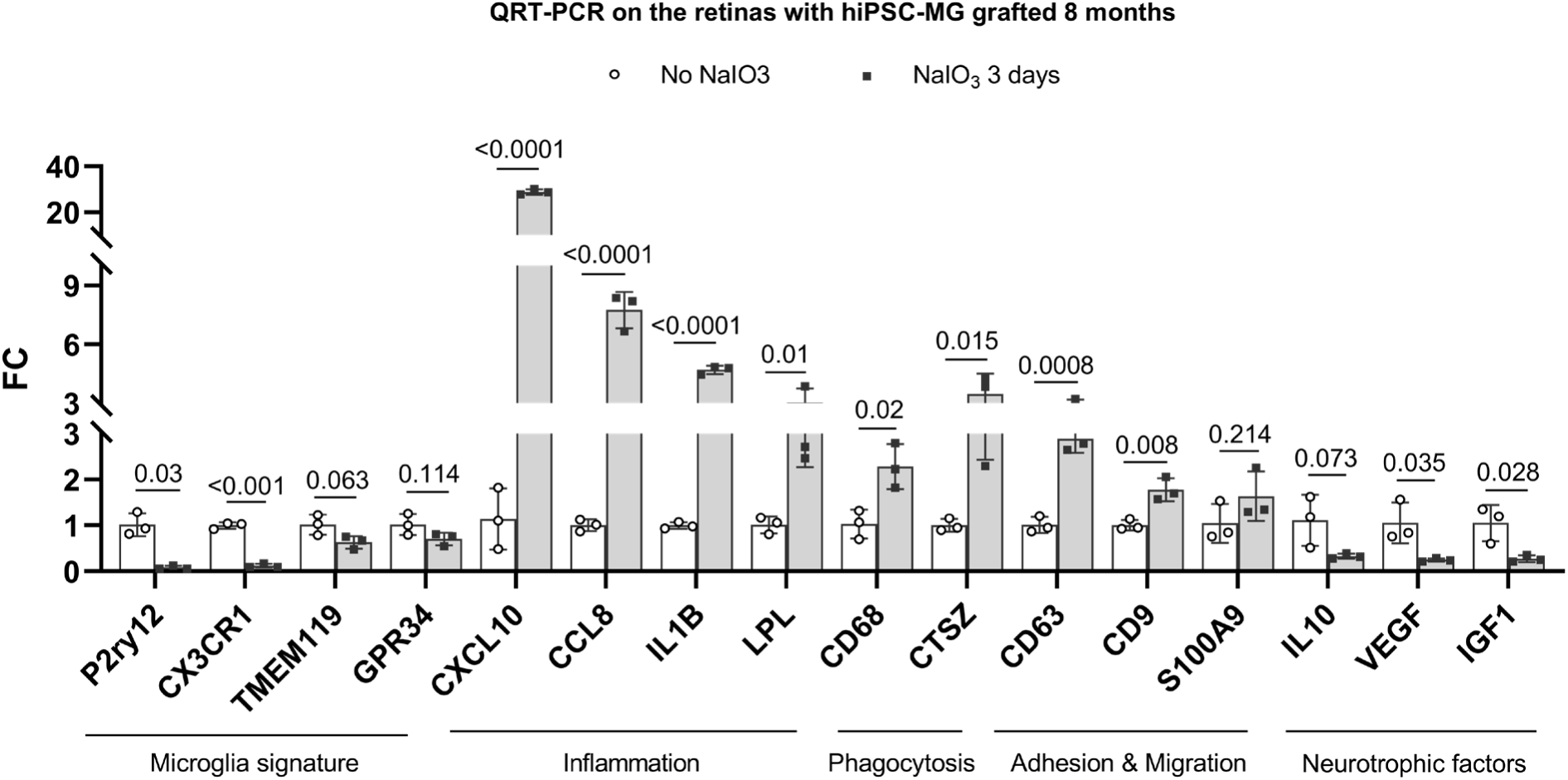
The inflammation, phagocytosis, adhesion and migration, neurotrophic factors, and microglia signature gene expression in hiPSC-derived microglia cells of grafted retinas. We compared gene coding sequences between humans and mice, chose a human-specific sequence to make the oligos (Suppl. Table 5), and ran a qRT-PCR on 8-month hiPSC-derived microglia cells grafted retinas with/without NaIO3-treated retinas. The results revealed that hiPSC-derived microglia cells expressed more inflammatory factors and phagocytosis genes and promoted cell migration but decreased microglia cell signature genes and neurotrophic factors in NaIO3-treated retina.

**Discussion suppl A.**
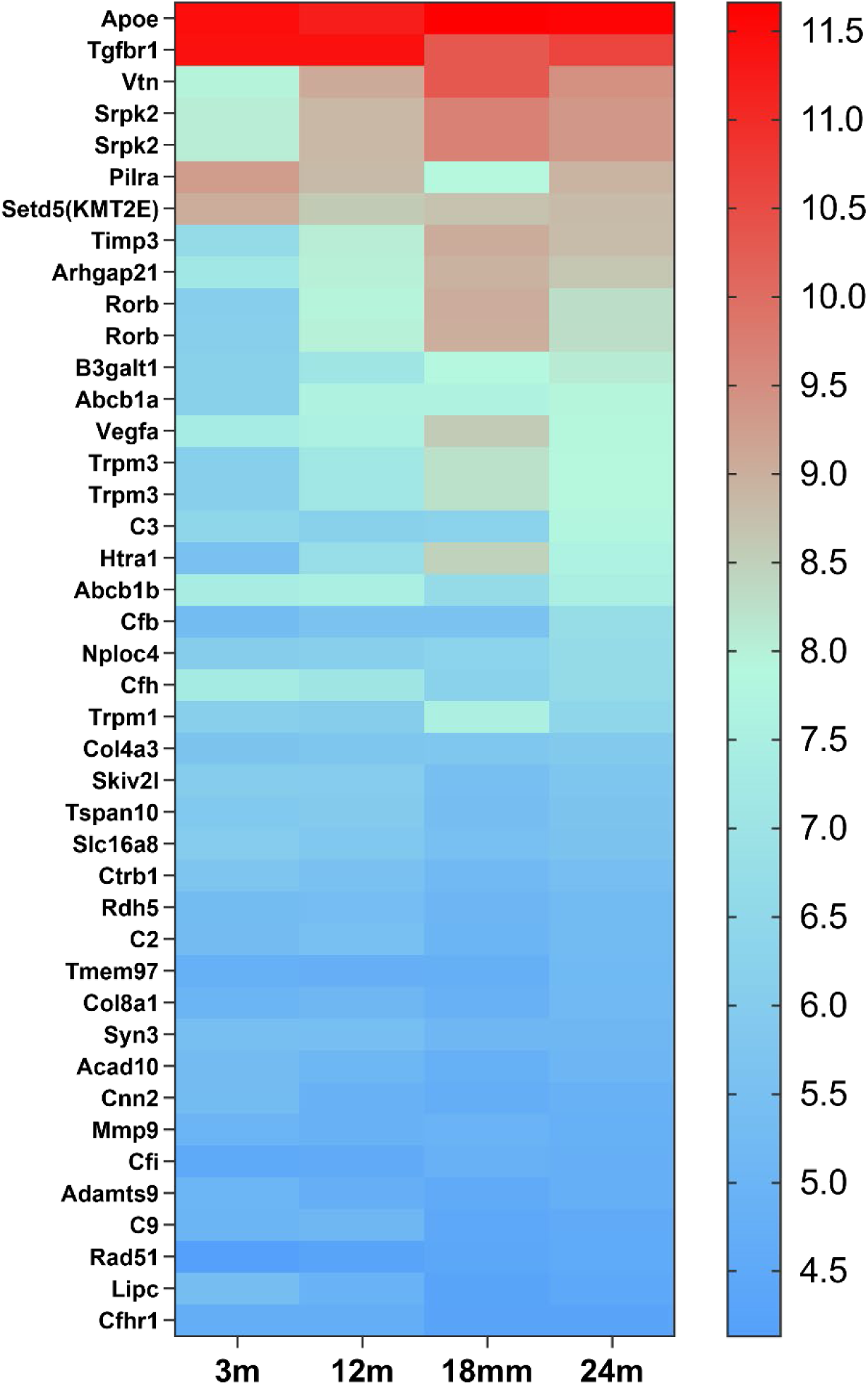
The heat map of 42 candidate genes from 34 loci associated with AMD expressed in retinal microglia cells. The microglia gene expression data are from microarray data previously published (Ma et al., 2013). The candidate genes came from the published paper (Den et al., 2022). The gene list is in Supplement Table 6.

**Discussion suppl B.**
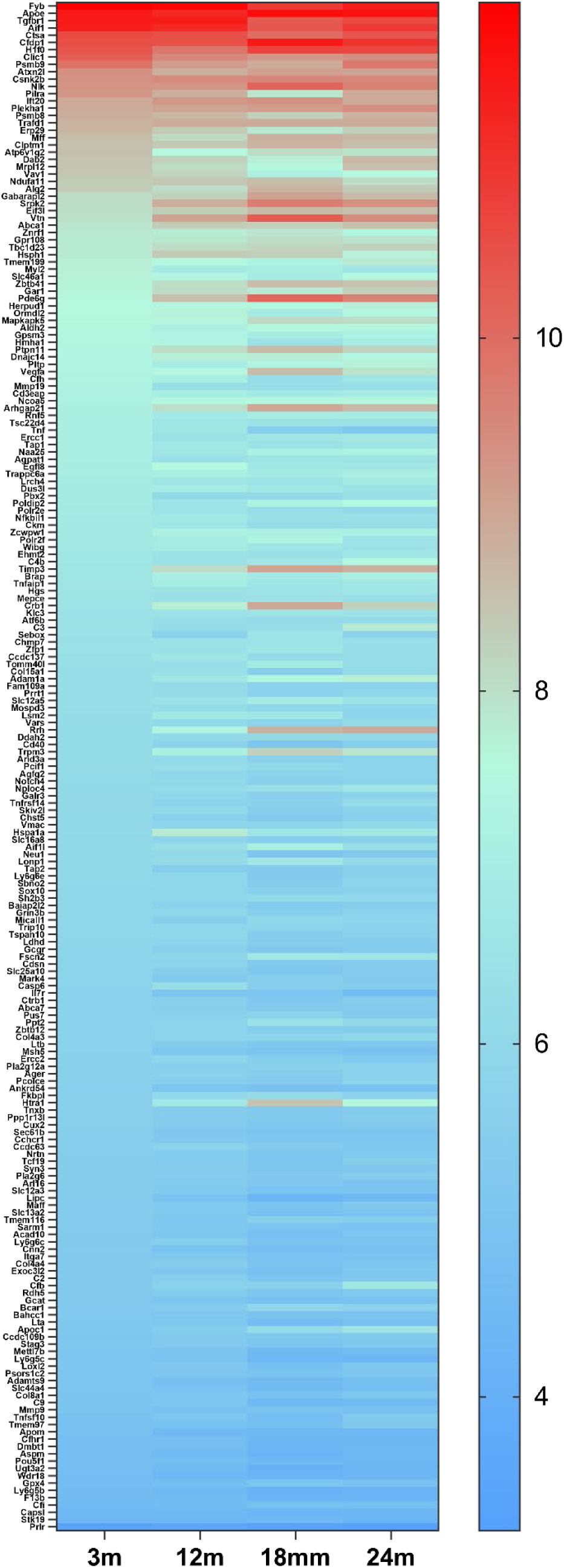
The heat map of 209 genes associated with AMD (Fritsche et al., 2016) expressed in retinal microglia cells. The microglia gene expression data are from microarray data previously published (Ma et al.,2013). The gene list is in Supplement Table 7.

## References

Abud EM, Ramirez RN, Martinez ES, Healy LM, Nguyen CHH, Newman SA, Yeromin AV, Scarfone VM, Marsh SE, Fimbres C, Caraway CA, Fote GM, Madany AM, Agrawal A, Kayed R, Gylys KH, Cahalan MD, Cummings BJ, Antel JP, Mortazavi A, Carson MJ, Poon WW, Blurton-Jones M. iPSC-Derived Human Microglia-like Cells to Study Neurological Diseases. Neuron. 2017 Apr 19;94(2):278–293.e9. doi: 10.1016/j.neuron.2017.03.042. PMID: 28426964; PMCID: PMC5482419.

Anderson SR, Roberts JM, Zhang J, Steele MR, Romero CO, Bosco A, et al. Developmental apoptosis promotes a disease-related gene signature and Independence from CSF1R signaling in retinal microglia. Cell Rep. 2019; 27(7):2002–13 e5.

Au NPB, Ma CHE. Neuroinflammation, Microglia and Implications for Retinal Ganglion Cell Survival and Axon Regeneration in Traumatic Optic Neuropathy. Front Immunol. 2022 Mar 4;13:860070. doi: 10.3389/fimmu.2022.860070. PMID: 35309305; PMCID: PMC8931466.

Beutner C, Lepperhof V, Dann A, Linnartz-Gerlach B, Litwak S, Napoli I, Prinz M, Neumann H. Engineered stem cell-derived microglia as therapeutic vehicle for experimental autoimmune encephalomyelitis. Gene Ther. 2013 Aug;20(8):797–806. doi: 10.1038/gt.2012.100. Epub 2013 Jan 17. PMID: 23324824.

Bohlen CJ, Bennett FC, Tucker AF, Collins HY, Mulinyawe SB, & Barres BA (2017). Diverse requirements for microglial survival, specification, and function revealed by defined-medium cultures. Neuron, 94(4), 759–773. 10.1016/j.neuron.2017.04.043 [PubMed: 28521131]

Bosco, A., Anderson, S. R., Roberts, J. M., Romero, C. O., Steele, M. R., and Vetter, M. L. (2019). Retinal microglia acquire a disease-associated transcriptome in chronic mouse glaucoma, which intensifies with neuroprotective complement inhibition. Invest. Ophthalmol. Vis. Sci. 60, 4002– 4002.

Böttcher C, Schlickeiser S, Sneeboer MAM, Kunkel D, Knop A, Paza E, Fidzinski P, Kraus L, Snijders GJL, Kahn RS, Schulz AR, Mei HE; NBB-Psy; Hol EM, Siegmund B, Glauben R, Spruth EJ, de Witte LD, Priller J. Human microglia regional heterogeneity and phenotypes determined by multiplexed single-cell mass cytometry. Nat Neurosci. 2019 Jan;22(1):78–90. doi: 10.1038/s41593-018-0290-2. Epub 2018 Dec 17. PMID: 30559476.

Broderick C, Hoek RM, Forrester JV, Liversidge J, Sedgwick JD, Dick AD. Constitutive retinal CD200 expression regulates resident microglia and activation state of inflammatory cells during experimental autoimmune uveoretinitis. Am J Pathol. 2002 Nov;161(5):1669–77. doi: 10.1016/S0002-9440(10)64444-6. PMID: 12414514; PMCID: PMC1850781.

Burns TC, Li MD, Mehta S, Awad AJ, & Morgan AA (2015). Mouse models rarely mimic the transcriptome of human neurodegenerative diseases: A systematic bioinformatics-based critique of preclinical models. European Journal of Pharmacology, 759, 101–117. 10.1016/j.ejphar.2015.03.021 [PubMed: 25814260].

Butovsky O, Jedrychowski MP, Moore CS, Cialic R, Lanser AJ, Gabriely G, Koeglsperger T, Dake B, Wu PM, Doykan CE, Fanek Z, Liu L, Chen Z, Rothstein JD, Ransohoff RM, Gygi SP, Antel JP, Weiner HL. Identification of a unique TGF-β-dependent molecular and functional signature in microglia. Nat Neurosci. 2014 Jan;17(1):131–43. doi: 10.1038/nn.3599. Epub 2013 Dec 8. Erratum in: Nat Neurosci. 2014 Sep;17(9):1286. PMID: 24316888; PMCID: PMC4066672.

Chadarevian JP, Lombroso SI, Peet GC, Hasselmann J, Tu C, Marzan DE, Capocchi J, Purnell FS, Nemec KM, Lahian A, Escobar A, England W, Chaluvadi S, O’Brien CA, Yaqoob F, Aisenberg WH, Porras-Paniagua M, Bennett ML, Davtyan H, Spitale RC, Blurton-Jones M, Bennett FC. Engineering an inhibitor-resistant human CSF1R variant for microglia replacement. J Exp Med. 2023 Mar 6;220(3):e20220857. doi: 10.1084/jem.20220857. Epub 2022 Dec 30. PMID: 36584406; PMCID: PMC9814156.

Checchin D, Sennlaub F, Levavasseur E, Leduc M, Chemtob S. Potential role of microglia in retinal blood vessel formation. Invest Ophthalmol Vis Sci. 2006;47(8):3595–602.

Colonna M, Butovsky O. Microglia function in the central nervous system during health and Neurodegeneration. Annu Rev Immunol. 2017;35:441–68. 4.

Combadière, C., Feumi, C., Raoul, W., Keller, N., Rodéro, M., Pézard, A., et al. (2007). CX3CR1-dependent subretinal microglia cell accumulation is associated with cardinal features of age-related macular degeneration. J. Clin. Invest. 117, 2920–2928. doi: 10.1172/JCI31692.

Cukras CA, Petrou P, Chew EY, Meyerle CB, Wong WT. Oral minocycline for the treatment of diabetic macular edema (DME): results of a phase I/II clinical study. Invest Ophthalmol Vis Sci. 2012 Jun 22;53(7):3865–74. doi: 10.1167/iovs.11-9413. PMID: 22589436; PMCID: PMC3390218.

Dagher NN, Najafi AR, Kayala KM, Elmore MR, White TE, Medeiros R, West BL, Green KN. Colony-stimulating factor 1 receptor inhibition prevents microglial plaque association and improves cognition in 3xTg-AD mice. J Neuroinflammation. 2015 Aug 1;12:139. doi: 10.1186/s12974-015-0366-9. PMID: 26232154; PMCID: PMC4522109.

Dawson TM, Golde TE, & Lagier-Tourenne C (2018). Animal models of neurodegenerative diseases. Nature Neuroscience, 21(10), 1370–1379. 10.1038/s41593-018-0236-8 [PubMed: 30250265]

Den Hollander AI, Mullins RF, Orozco LD, Voigt AP, Chen HH, Strunz T, Grassmann F, Haines JL, Kuiper JJW, Tumminia SJ, Allikmets R, Hageman GS, Stambolian D, Klaver CCW, Boeke JD, Chen H, Honigberg L, Katti S, Frazer KA, Weber BHF, Gorin MB. Systems genomics in age-related macular degeneration. Exp Eye Res. 2022 Dec;225:109248. doi: 10.1016/j.exer.2022.109248. Epub 2022 Sep 13. PMID: 36108770; PMCID: PMC10150562.

Douvaras P, Sun B, Wang M, Kruglikov I, Lallos G, Zimmer M, Terrenoire C, Zhang B, Gandy S, Schadt E, Freytes DO, Noggle S, Fossati V. Directed Differentiation of Human Pluripotent Stem Cells to Microglia. Stem Cell Reports. 2017 Jun 6;8(6):1516–1524. doi: 10.1016/j.stemcr.2017.04.023. Epub 2017 May 18. PMID: 28528700; PMCID: PMC5470097.

Eisenberg E, Levanon EY. Human housekeeping genes, revisited. Trends Genet. 2013 Oct;29(10):569–74. doi: 10.1016/j.tig.2013.05.010. Epub 2013 Jun 27. Erratum in: Trends Genet. 2014 Mar;30(3):119-20. PMID: 23810203.

Elmore MRP, Hohsfield LA, Kramár EA, Soreq L, Lee RJ, Pham ST, Najafi AR, Spangenberg EE, Wood MA, West BL, Green KN. Replacement of microglia in the aged brain reverses cognitive, synaptic, and neuronal deficits in mice. Aging Cell. 2018 Dec;17(6):e12832. doi: 10.1111/acel.12832. Epub 2018 Oct 2. PMID: 30276955; PMCID: PMC6260908.

Friedman BA, Srinivasan K, Ayalon G, Meilandt WJ, Lin H, Huntley MA, Cao Y, Lee SH, Haddick PCG, Ngu H, Modrusan Z, Larson JL, Kaminker JS, van der Brug MP, Hansen DV. Diverse Brain Myeloid Expression Profiles Reveal Distinct Microglial Activation States and Aspects of Alzheimer’s Disease Not Evident in Mouse Models. Cell Rep. 2018 Jan 16;22(3):832–847. doi: 10.1016/j.celrep.2017.12.066. PMID: 29346778.

Fritsche LG, Igl W, Bailey JN, Grassmann F, … Weber BH, Abecasis GR, Heid IM. A large genome-wide association study of age-related macular degeneration highlights contributions of rare and common variants. Nat Genet. 2016 Feb;48(2):134–43. doi: 10.1038/ng.3448. Epub 2015 Dec 21. PMID: 26691988; PMCID: PMC4745342.

Galatro TF, Holtman IR, Lerario AM, Vainchtein ID, Brouwer N, Sola PR, Veras MM, Pereira TF, Leite REP, Möller T, Wes PD, Sogayar MC, Laman JD, den Dunnen W, Pasqualucci CA, Oba-Shinjo SM, Boddeke EWGM, Marie SKN, Eggen BJL. Transcriptomic analysis of purified human cortical microglia reveals age-associated changes. Nat Neurosci. 2017 Aug;20(8):1162–1171. doi: 10.1038/nn.4597. Epub 2017 Jul 3. PMID: 28671693.

Gosselin D, Skola D, Coufal NG, Holtman IR, Schlachetzki JCM, Sajti E, Jaeger BN, O’Connor C, Fitzpatrick C, Pasillas MP, Pena M, Adair A, Gonda DD, Levy ML, Ransohoff RM, Gage FH, Glass CK. An environment-dependent transcriptional network specifies human microglia identity. Science. 2017 Jun 23;356(6344):eaal3222. doi: 10.1126/science.aal3222. Epub 2017 May 25. PMID: 28546318; PMCID: PMC5858585.

Green, D. R., Oguin, T. H., and Martinez, J. (2016). The clearance of dying cells: table for two. Cell Death Differ. 23, 915–926. doi: 10.1038/cdd.2015.172.

Guilliams M, Scott CL. Does niche competition determine the origin of tissue-resident macrophages? Nat Rev Immunol. 2017 Jul;17(7):451–460. doi: 10.1038/nri.2017.42. Epub 2017 May 2. PMID: 28461703.

Guo S, Cázarez-Márquez F, Jiao H, Foppen E, Korpel NL, Grootemaat AE, Liv N, Gao Y, van der Wel N, Zhou B, Nie G, Yi CX. Specific Silencing of Microglial Gene Expression in the Rat Brain by Nanoparticle-Based Small Interfering RNA Delivery. ACS Appl Mater Interfaces. 2022 Feb 2;14(4):5066–5079. doi: 10.1021/acsami.1c22434. Epub 2022 Jan 18. PMID: 35041392; PMCID: PMC8815040.

Haenseler W, Sansom SN, Buchrieser J, Newey SE, Moore CS, Nicholls FJ, Chintawar S, Schnell C, Antel JP, Allen ND, Cader MZ, Wade-Martins R, James WS, Cowley SA. A Highly Efficient Human Pluripotent Stem Cell Microglia Model Displays a Neuronal-Co-culture-Specific Expression Profile and Inflammatory Response. Stem Cell Reports. 2017 Jun 6;8(6):1727–1742. doi: 10.1016/j.stemcr.2017.05.017. PMID: 28591653; PMCID: PMC5470330.

Han J, Sarlus H, Wszolek ZK, Karrenbauer VD, Harris RA. Microglial replacement therapy: a potential therapeutic strategy for incurable CSF1R-related leukoencephalopathy. Acta Neuropathol Commun. 2020 Dec 7;8(1):217. doi: 10.1186/s40478-020-01093-3. PMID: 33287883; PMCID: PMC7720517.

Hasselmann J, Blurton-Jones M. Human iPSC-derived microglia: A growing toolset to study the brain’s innate immune cells. Glia. 2020 Apr;68(4):721–739. doi: 10.1002/glia.23781. Epub 2020 Jan 11. PMID: 31926038; PMCID: PMC7813153.

Hohsfield LA, Najafi AR, Ghorbanian Y, Soni N, Hingco EE, Kim SJ, Jue AD, Swarup V, Inlay MA, Green KN. Effects of long-term and brain-wide colonization of peripheral bone marrow-derived myeloid cells in the CNS. J Neuroinflammation. 2020 Sep 20;17(1):279. doi: 10.1186/s12974-020-01931-0. PMID: 32951604; PMCID: PMC7504855.

Hong S, Dissing-Olesen L, Stevens B. New insights on the role of microglia in synaptic pruning in health and disease. Curr Opin Neurobiol. 2016;36: 128 –34.

Huang T, Cui J, Li L, Hitchcock PF, Li Y. The role of microglia in the neurogenesis of zebrafish retina. Biochem Biophys Res Commun. 2012; 421(2):214 –20

Huang Y, Xu Z, Xiong S, Qin G, Sun F, Yang J, Yuan TF, Zhao L, Wang K, Liang YX, Fu L, Wu T, So KF, Rao Y, Peng B. Dual extra-retinal origins of microglia in the model of retinal microglia repopulation. Cell Discov. 2018 Feb 27;4:9. doi: 10.1038/s41421-018-0011-8. PMID: 29507754; PMCID: PMC5827656.

Karlsson KR, Cowley S, Martinez FO, Shaw M, Minger SL, James W. Homogeneous monocytes and macrophages from human embryonic stem cells following coculture-free differentiation in M-CSF and IL-3. Exp Hematol. 2008 Sep;36(9):1167–75. doi: 10.1016/j.exphem.2008.04.009. Epub 2008 Jun 11. PMID: 18550257; PMCID: PMC2635571.

Karlstetter M, Scholz R, Rutar M, Wong WT, Provis JM, Langmann T. Retinal microglia: just bystander or target for therapy? Prog Retin Eye Res. 2015 Mar;45:30–57. doi: 10.1016/j.preteyeres.2014.11.004. Epub 2014 Dec 2. PMID: 25476242.

Kubota Y, Takubo K, Shimizu T, Ohno H, Kishi K, Shibuya M, et al. M-CSF inhibition selectively targets pathological angiogenesis and lymphangiogenesis. J Exp Med. 2009;206(5):1089–102

Larochelle A, Bellavance MA, Michaud JP, Rivest S. Bone marrow-derived macrophages and the CNS: An update on the use of experimental chimeric mouse models and bone marrow transplantation in neurological disorders. Biochim Biophys Acta. 2016 Mar;1862(3):310–22. doi: 10.1016/j.bbadis.2015.09.017. Epub 2015 Oct 8. PMID: 26432480.

Leach LL, Croze RH, Hu Q, Nadar VP, Clevenger TN, Pennington BO, Gamm DM, Clegg DO. Induced Pluripotent Stem Cell-Derived Retinal Pigmented Epithelium: A Comparative Study Between Cell Lines and Differentiation Methods. J Ocul Pharmacol Ther. 2016 Jun;32(5):317–30. doi: 10.1089/jop.2016.0022. Epub 2016 May 16. PMID: 27182743; PMCID: PMC5911695.

Lee JE, Liang KJ, Fariss RN, Wong WT. Ex vivo dynamic imaging of retinal microglia using time-lapse confocal microscopy. Invest Ophthalmol Vis Sci. 2008 Sep;49(9):4169–76. doi: 10.1167/iovs.08-2076. Epub 2008 May 16. PMID: 18487378; PMCID: PMC2652634.

Li Q, Barres BA. Microglia and macrophages in brain homeostasis and disease. Nat Rev Immunol. 2018;18(4):225–42. 3.

Lund H, Pieber M, Parsa R, Han J, Grommisch D, Ewing E, Kular L, Needhamsen M, Espinosa A, Nilsson E, Överby AK, Butovsky O, Jagodic M, Zhang XM, Harris RA. Competitive repopulation of an empty microglial niche yields functionally distinct subsets of microglia-like cells. Nat Commun. 2018 Nov 19;9(1):4845. doi: 10.1038/s41467-018-07295-7. PMID: 30451869; PMCID: PMC6242869.

Ma W, Cojocaru R, Gotoh N, Gieser L, Villasmil R, Cogliati T, Swaroop A, Wong WT. Gene expression changes in aging retinal microglia: relationship to microglial support functions and regulation of activation. Neurobiol Aging. 2013 Oct;34(10):2310–21. doi: 10.1016/j.neurobiolaging.2013.03.022. Epub 2013 Apr 19. PMID: 23608111; PMCID: PMC3706521.

Ma W, Zhang Y, Gao C, Fariss RN, Tam J, Wong WT. Monocyte infiltration and proliferation reestablish myeloid cell homeostasis in the mouse retina following retinal pigment epithelial cell injury. Sci Rep. 2017 Aug 16;7(1):8433. doi: 10.1038/s41598-017-08702-7. PMID: 28814744; PMCID: PMC5559448.

Ma W, Zhao L, Fontainhas AM, Fariss RN, Wong WT. Microglia in the mouse retina alter the structure and function of retinal pigmented epithelial cells: a potential cellular interaction relevant to AMD. PLoS One. 2009 Nov 20;4(11):e7945. doi: 10.1371/journal.pone.0007945. PMID: 19936204; PMCID: PMC2775955.

Ma W, Zhao L, Wong WT. Microglia in the outer retina and their relevance to pathogenesis of age-related macular degeneration. Adv Exp Med Biol. 2012;723:37–42. doi:10.1007/978-1-4614-0631-0_6

Mancuso R, Van Den Daele J, Fattorelli N, Wolfs L, Balusu S, Burton O, Liston A, Sierksma A, Fourne Y, Poovathingal S, Arranz-Mendiguren A, Sala Frigerio C, Claes C, Serneels L, Theys T, Perry VH, Verfaillie C, Fiers M, De Strooper B. Stem-cell-derived human microglia transplanted in mouse brain to study human disease. Nat Neurosci. 2019 Dec;22(12):2111–2116. doi: 10.1038/s41593-019-0525-x. Epub 2019 Oct 28. PMID: 31659342.

Muffat J, Li Y, Yuan B, Mitalipova M, Omer A, Corcoran S, Bakiasi G, Tsai LH, Aubourg P, Ransohoff RM, Jaenisch R. Efficient derivation of microglia-like cells from human pluripotent stem cells. Nat Med. 2016 Nov;22(11):1358–1367. doi: 10.1038/nm.4189. Epub 2016 Sep 26. PMID: 27668937; PMCID: PMC5101156.

Nau R, Ribes S, Djukic M, Eiffert H. Strategies to increase the activity of microglia as efficient protectors of the brain against infections. Front Cell Neurosci. 2014 May 22;8:138. doi: 10.3389/fncel.2014.00138. PMID: 24904283; PMCID: PMC4033068.

Neumann H. Microglia: a cellular vehicle for CNS gene therapy. J Clin Invest. 2006 Nov;116(11):2857–60. doi: 10.1172/JCI30230. PMID: 17080190; PMCID: PMC1626126.

Okunuki Y, Mukai R, Nakao T, Tabor SJ, Butovsky O, Dana R, Ksander DR, Connor KM. Retinal microglia initiate neuroinflammation in ocular autoimmunity. PNAS, 2019, 116 (20) 9989–9998; DOI: 10.1073/pnas.1820387116

Pandya H, Shen MJ, Ichikawa DM, Sedlock AB, Choi Y, Johnson KR, Kim G, Brown MA, Elkahloun AG, Maric D, Sweeney CL, Gossa S, Malech HL, McGavern DB, Park JK. Differentiation of human and murine induced pluripotent stem cells to microglia-like cells. Nat Neurosci. 2017 May;20(5):753–759. doi: 10.1038/nn.4534. Epub 2017 Mar 2. PMID: 28253233; PMCID: PMC5404968.

Paolicelli RC, Bolasco G, Pagani F, Maggi L, Scianni M, Panzanelli P, Giustetto M, Ferreira TA, Guiducci E, Dumas L, Ragozzino D, Gross CT. Synaptic pruning by microglia is necessary for normal brain development. Science. 2011 Sep 9;333(6048):1456-8. doi: 10.1126/science.1202529. Epub 2011 Jul 21. PMID: 21778362.

Parajuli B, Saito H, Shinozaki Y, Shigetomi E, Miwa H, Yoneda S, Tanimura M, Omachi S, Asaki T, Takahashi K, Fujita M, Nakashima K, Koizumi S. Transnasal transplantation of human induced pluripotent stem cell-derived microglia to the brain of immunocompetent mice. Glia. 2021 Oct;69(10):2332–2348. doi: 10.1002/glia.23985. Epub 2021 Jul 26. PMID: 34309082.

Pocock JM, Piers TM. Modelling microglial function with induced pluripotent stem cells: an update. Nat Rev Neurosci. 2018 Aug;19(8):445–452. doi: 10.1038/s41583-018-0030-3. PMID: 29977068.

Puñal V, Paisley C, Brecha F, Lee M, Perelli R, Wang J et al. Large-scale death of retinal astrocytes during normal development is non-apoptotic and implemented by microglia. PLoS Biology, 2019:17(10).

Ramírez AI, de Hoz R, Fernández-Albarral JA, Salobrar-Garcia E, Rojas B, Valiente-Soriano FJ, Avilés-Trigueros M, Villegas-Pérez MP, Vidal-Sanz M, Triviño A, Ramírez JM, Salazar JJ. Time course of bilateral microglial activation in a mouse model of laser-induced glaucoma. Sci Rep. 2020 Mar 17;10(1):4890. doi: 10.1038/s41598-020-61848-9. PMID: 32184450; PMCID: PMC7078298.

Réu P, Khosravi A, Bernard S, Mold JE, Salehpour M, Alkass K, Perl S, Tisdale J, Possnert G, Druid H, Frisén J. The Lifespan and Turnover of Microglia in the Human Brain. Cell Rep. 2017 Jul 25;20(4):779–784. doi: 10.1016/j.celrep.2017.07.004. PMID: 28746864; PMCID: PMC5540680.

Ritter MR, Banin E, Moreno SK, Aguilar E, Dorrell MI, Friedlander M. Myeloid progenitors differentiate into microglia and promote vascular repair in a model of ischemic retinopathy. J Clin Invest. 2006;116(12):3266–76. 104.

Schafer DP, Lehrman EK, Kautzman AG, Koyama R, Mardinly AR, Yamasaki R, et al. Microglia sculpt postnatal neural circuits in an activity and complement-dependent manner. Neuron. 2012;74(4):691–705.

Schafer DP, Stevens B. Microglia function in central nervous system development and plasticity. Cold Spring Harb Perspect Biol. 2015;7(10): a020545. 133.

Shi Q, Chang C, SalIba1 A, Bhat MA. Microglial mTOR Activation Upregulates Trem2 and Enhances β-Amyloid Plaque Clearance in the 5XFAD Alzheimer’s Disease Model. J Neurosci. 2022 Jul 6;42(27):5294-5313. doi: 10.1523/JNEUROSCI.2427-21.2022. Epub 2022 Jun 7. PMID: 35672148; PMCID: PMC9270922.

Shibuya Y, Kumar KK, Mader MM, Yoo Y, Ayala LA, Zhou M, Mohr MA, Neumayer G, Kumar I, Yamamoto R, Marcoux P, Liou B, Bennett FC, Nakauchi H, Sun Y, Chen X, Heppner FL, Wyss-Coray T, Südhof TC, Wernig M. Treatment of a genetic brain disease by CNS-wide microglia replacement. Sci Transl Med. 2022 Mar 16;14(636):eabl9945. doi: 10.1126/scitranslmed.abl9945. Epub 2022 Mar 16. PMID: 35294256; PMCID: PMC9618306.

Sierra A, Abiega O, Shahraz A, Neumann H. Janus-faced microglia: beneficial and detrimental consequences of microglial phagocytosis. Front Cell Neurosci. 2013;7:6.

Silverman SM, Ma W, Wang X, Zhao L, Wong WT. C3- and CR3-dependent microglial clearance protects photoreceptors in retinitis pigmentosa. J Exp Med. 2019 Aug 5;216(8):1925–1943. doi: 10.1084/jem.20190009. Epub 2019 Jun 17. PMID: 31209071; PMCID: PMC6683998.

Smith AM, & Dragunow M (2014). The human side of microglia. Trends in Neurosciences, 37(3), 125–135. 10.1016/j.tins.2013.12.001 [PubMed: 24388427]

Stevens B, Allen NJ, Vazquez LE, Howell GR, Christopherson KS, Nouri N, et al. The classical complement cascade mediates CNS synapse elimination. Cell. 2007;131(6):1164–78

Svoboda DS, Barrasa MI, Shu J, Rietjens R, Zhang S, Mitalipova M, Berube P, Fu D, Shultz LD, Bell GW, Jaenisch R. Human iPSC-derived microglia assume a primary microglia-like state after transplantation into the neonatal mouse brain. Proc Natl Acad Sci U S A. 2019 Dec 10;116(50):25293–25303. doi: 10.1073/pnas.1913541116. Epub 2019 Nov 26. PMID: 31772018; PMCID: PMC6911218.

Takata K, Kozaki T, Lee CZW, Thion MS, Otsuka M, Lim S, Utami KH, Fidan K, Park DS, Malleret B, Chakarov S, See P, Low D, Low G, Garcia-Miralles M, Zeng R, Zhang J, Goh CC, Gul A, Hubert S, Lee B, Chen J, Low I, Shadan NB, Lum J, Wei TS, Mok E, Kawanishi S, Kitamura Y, Larbi A, Poidinger M, Renia L, Ng LG, Wolf Y, Jung S, Önder T, Newell E, Huber T, Ashihara E, Garel S, Pouladi MA, Ginhoux F. Induced-Pluripotent-Stem-Cell-Derived Primitive Macrophages Provide a Platform for Modeling Tissue-Resident Macrophage Differentiation and Function. Immunity. 2017 Jul 18;47(1):183–198.e6. doi: 10.1016/j.immuni.2017.06.017. Erratum in: Immunity. 2020 Feb 18;52(2):417-418. PMID: 28723550.

Tanaka T, Yokoi T, Tamalu F, Watanabe S, Nishina S, Azuma N. Generation of Retinal Ganglion Cells With Functional Axons From Mouse Embryonic Stem Cells and Induced Pluripotent Stem Cells. Invest Ophthalmol Vis Sci. 2016 Jun 1;57(7):3348–59. doi: 10.1167/iovs.16-19166. PMID: 27367502.

Ueda Y, Gullipalli D, & Song WC (2016). Modeling complement-driven diseases in transgenic mice: Values and limitations. Immunobiology, 221(10), 1080–1090. 10.1016/j.imbio.2016.06.007 [PubMed: 27371974]

Ulland TK, Colonna M. TREM2 - a key player in microglial biology and Alzheimer disease. Nat Rev Neurol. 2018 Nov;14(11):667–675. doi: 10.1038/s41582-018-0072-1. PMID: 30266932.

van der Poel M, Ulas T, Mizee MR, Hsiao CC, Miedema SSM, Adelia, Schuurman KG, Helder B, Tas SW, Schultze JL, Hamann J, Huitinga I. Transcriptional profiling of human microglia reveals grey-white matter heterogeneity and multiple sclerosis-associated changes. Nat Commun. 2019 Mar 13;10(1):1139. doi: 10.1038/s41467-019-08976-7. PMID: 30867424; PMCID: PMC6416318.

Van Wilgenburg B, Browne C, Vowles J, Cowley SA. Efficient, long term production of monocyte-derived macrophages from human pluripotent stem cells under partly-defined and fully-defined conditions. PLoS One. 2013 Aug 12;8(8):e71098. doi: 10.1371/journal.pone.0071098. PMID: 23951090; PMCID: PMC3741356.

Wang M, Wang X, Zhao L, Ma W, Rodriguez IR, Fariss RN, Wong WT. Macroglia-microglia interactions via TSPO signaling regulates microglial activation in the mouse retina. J Neurosci. 2014 Mar 5;34(10):3793–806. doi: 10.1523/JNEUROSCI.3153-13.2014. PMID: 24599476; PMCID: PMC3942591.

Wang S, Mustafa M, Yuede CM, Salazar SV, Kong P, Long H, Ward M, Siddiqui O, Paul R, Gilfillan S, Ibrahim A, Rhinn H, Tassi I, Rosenthal A, Schwabe T, Colonna M. Anti-human TREM2 induces microglia proliferation and reduces pathology in an Alzheimer’s disease model. J Exp Med. 2020 Sep 7;217(9):e20200785. doi: 10.1084/jem.20200785. PMID: 32579671; PMCID: PMC7478730.

Wang X, Zhao L, Zhang J, Fariss RN, Ma W, Kretschmer F, Wang M, Qian HH, Badea TC, Diamond JS, Gan WB, Roger JE, Wong WT. Requirement for Microglia for the Maintenance of Synaptic Function and Integrity in the Mature Retina. J Neurosci. 2016 Mar 2;36(9):2827–42. doi: 10.1523/JNEUROSCI.3575-15.2016. PMID: 26937019; PMCID: PMC4879218.

Willis EF, MacDonald KPA, Nguyen QH, Garrido AL, Gillespie ER, Harley SBR, Bartlett PF, Schroder WA, Yates AG, Anthony DC, Rose-John S, Ruitenberg MJ, Vukovic J. Repopulating Microglia Promote Brain Repair in an IL-6-Dependent Manner. Cell. 2020 Mar 5;180(5):833–846.e16. doi: 10.1016/j.cell.2020.02.013. PMID: 32142677.

Xu H, Chen M, Diabetic retinopathy and dysregulated innate immunity, Vision Research, Volume 139, 2017, Pages 39-46, ISSN 0042-6989.

Xu R, Li X, Boreland AJ, Posyton A, Kwan K, Hart RP, Jiang P. Human iPSC-derived microgliaretain their identity and functionally integrate in the chimeric mouse brain. Nat Commun. 2020 Mar 27;11(1):1577. doi: 10.1038/s41467-020-15411-9. PMID: 32221280; PMCID: PMC7101330.

Xu Z, Rao Y, Huang Y, Zhou T, Feng R, Xiong S, Yuan TF, Qin S, Lu Y, Zhou X, Li X, Qin B, Mao Y, Peng B. Efficient Strategies for Microglia Replacement in the Central Nervous System. Cell Rep. 2020 Nov 24;33(8):108443. doi: 10.1016/j.celrep.2020.108443. Erratum for: Cell Rep. 2020 Aug 11;32(6):108041. PMID: 33238120.

Zhang Y, Zhao L, Wang X, Ma W, Lazere A, Qian HH, Zhang J, Abu-Asab M, Fariss RN, Roger JE, Wong WT. Repopulating retinal microglia restore endogenous organization and function under CX3CL1-CX3CR1 regulation. Sci Adv. 2018 Mar 21;4(3):eaap8492. doi: 10.1126/sciadv.aap8492. PMID: 29750189; PMCID: PMC5943055.

Zhao L, Ma W, Fariss RN, Wong WT. Minocycline attenuates photoreceptor degeneration in a mouse model of subretinal hemorrhage microglial: inhibition as a potential therapeutic strategy. Am J Pathol. 2011 Sep;179(3):1265–77. doi: 10.1016/j.ajpath.2011.05.042. Epub 2011 Jul 19. PMID: 21763674; PMCID: PMC3157282.

Zhao L, Zabel MK, Wang X, Ma W, Shah P, Fariss RN, Qian H, Parkhurst CN, Gan WB, Wong WT. Microglial phagocytosis of living photoreceptors contributes to inherited retinal degeneration. EMBO Mol Med. 2015 Sep;7(9):1179–97. doi: 10.15252/emmm.201505298. PMID: 26139610; PMCID: PMC4568951.

Zhong X, Gutierrez C, Xue T, Hampton C, Vergara MN, Cao LH, Peters A, Park TS, Zambidis ET, Meyer JS, Gamm DM, Yau KW, Canto-Soler MV. Generation of three-dimensional retinal tissue with functional photoreceptors from human iPSCs. Nat Commun. 2014 Jun 10;5:4047. doi: 10.1038/ncomms5047. PMID: 24915161; PMCID: PMC4370190.

Zhou J, Yang J, Dai M, Lin D, Zhang R, Liu H, Yu A, Vakal S, Wang Y, Li X. A combination of inhibiting microglia activity and remodeling gut microenvironment suppresses the development and progression of experimental autoimmune uveitis. Biochem Pharmacol. 2020 Oct;180:114108. doi: 10.1016/j.bcp.2020.114108. Epub 2020 Jun 20. PMID: 32569628.

