## Supplement table 4 for "Human iPSC-derived Microglial Cells Integrated into Mouse Retina and Recapitulated Features of Endogenous Microglia"

**Supplement Table 1. Sequences of oligonucleotide primers used in polymerase chain reaction (PCR) assays**

| Name | Reverse (5’-3’) | Forward (5’-3’) |
| --- | --- | --- |
| Rps13 | GCGACCAGGGCAGAGGC | CTGATCTTCCTGAAGATCTCT |
| GAPDH | CTCCTTGGAGGCCATGTGG | CTACACTGAGCACCAGGTG |
| Il1a | CTACGCCTGGTTTTCCAGTA | CCAACCTCCTCTTCTTCTGG |
| IL1B | GGAAGACACAAATTGCATGGTG | CTCTACAGCTGGAGAGTGTAG |
| IL6 | CTACATTTGCCGAAGAGCCC | GTCCTGATCCAGTTCCTGCA |
| IL8 | GAATCCATCCCCCTGTTTTCA | GACATACTCCAAACCTTTCCAC |
| TNFa | CACAGGGCAATGATCCCAAAG | CCAAGGTCAACCTCCTCTCT |
| CXCL10 | GGAGATCTTTTAGACCTTTCC | CCTGTTAATCCAAGGTCTTTAG |
| MCP1 | CAAGTCTTCGGAGTTTGGGT | CTTCACCAATAGGAAGATCTC |
| CCL3 | CAGGCACTCAGCTCCAGG | CTGCTGCTTCAGCTACACC |
| CCL4 | CAGTTCAGTTCCAGGTCATAC | CTGCTTTTCTTACACCGCGAG |
| IL10 | GTTTCGTATCTTCATTGTCATGTA | CCCTCAGGCTGAGGCTACG |
| CX3CR1 | CAGAGAAGGAGCAATGCATC | CCTCTCATCTATGCATTTGCT |
| P2Ry12 | CATTGGAGTCTCTTCATTTGG | GAGAGCACTCTGTGGTTAAC |
| CD11b | CTACTGGGGTTCGGCCCC | GGTCCCAGACGGAGACCA |
| Iba1 | CACATTTTTAGGATGGCAGATC | CCATCCTCTGCCCCCAGAT |
| CTSS | CTAGATTTCTGGGTAAGAGGG | CTACTATGAACCATCCTGTAC |
| Hexb | CATGTTCTCATGGTTACAATATC | GTGGATGCAACTAACCTCACT |
| Sall1 | CTCGTGACGATCTCCTTGC | GGTGGCATCCCTCCAATTC |
| IL34 | CAGGGCAAGAGGCCCTCG | GCTCCTGCTGTAAACAAAGCT |
| C1qa | CAGGCAGATGGGAAGATGAG | GACATACTCCAAACCTTTCCAC |
| C1qb | CAGGCCTCCATATCTGGAAAG | GGAGCGTGCACAGAAGGTG |
| ApoE | CAGTGATTGTCGCTGGGCAC | GAAGGAGCAGGTGGCGGAG |
| CD45 | CTATGAACCTTGATTTAAAGCTG | GTCATTGCCAGCACCTACCC |
| CD68 | CAGAGGGCCTGGTAGGCG | GTCCACCTCGACCTGCTCT |
| CD74 | CACATGGGGACTGGGCCC | CATTCAGGCCCAAGTGCGAC |
| VEGF | CACCGCCTCGGCTTGTCAC | GCCCGCTGCTGTCTAATGC |
